# Structure of an active FlhA ring reveals allosteric coupling between substrate specificity and export engine activation

**DOI:** 10.64898/2026.05.29.728899

**Authors:** Miki Kinoshita, Tohru Minamino, Hiroyuki Terashima, Kazuki Kasai, Keiichi Namba

## Abstract

**The bacterial flagellar type III secretion system transports structural subunits in a defined order to assemble the flagellum. FlhA is the core component of this nanomachine; its transmembrane domain functions as an ion-driven export engine and its cytoplasmic domain connected by a flexible linker forms a nonameric ring that controls substrate selection. The coupling mechanism of these activities has remained unclear because the transmembrane domain has resisted structural analysis. Here we identify linker deletions that autonomously activate the export gate and capture a functionally active, hook-type protein export state of the entire ring complex. Using cryo-electron microscopy, we determine the structure of the FlhA(Δ331-347) ring at 2.73 Å resolution. The structure shows linker-mediated docking of the cytoplasmic ring to the transmembrane ring to stabilize the entire ring while sterically excluding filament-type substrate binding. These findings reveal how the FlhA ring allosterically couples substrate specificity to ion-driven export during flagellar assembly.**

---

The bacterial flagellum is a supramolecular motility machine consisting of the basal body, which functions as a rotary motor, the filament, which acts as a helical propeller, and the hook, which works as a universal joint connecting the basal body and filament. The flagellum is assembled in the cell exterior by exporting subunit proteins via the flagellar type III secretion system (fT3SS), a membrane-embedded nanomachine that exports structural subunits in a defined order (Extended Data Fig. 1). During hook assembly, the fT3SS selectively transports hook-type substrates (FlgD, FlgE, and FliK) and then switches to filament-type substrate export (FlgK, FlgL, FliC, FliD) once the hook reaches its mature length. This substrate specificity switching is essential for efficient and robust flagellar morphogenesis^1^.

The fT3SS comprises a transmembrane export gate complex powered by proton motive force across the cytoplasmic membrane and a cytoplasmic ATPase complex. The ATPase complex, composed of FliH, FliI and FliJ, functions as an ATP-driven activator that enables the export gate—comprising FlhA, FlhB, FliP, FliQ and FliR—to operate as an active H⁺-driven protein transporter^2,3^. In addition, the export gate is equipped with a membrane voltage sensor^4^ and can utilize Na⁺ as an alternative coupling ion^5,6^, allowing autonomous activation under conditions of elevated membrane potential or increased external Na⁺ concentration, even in the absence of the ATPase complex.

FlhA is the core component of the export gate that plays dual roles in energy transduction and substrate selection. Its N-terminal transmembrane domain (FlhA_TM_) conducts both H^+^ and Na^+^ to power protein export^5^, whereas its C-terminal cytoplasmic domain (FlhA_C_) forms a nonameric ring that serves as a docking platform for the ATPase component FliJ and the flagellar export chaperones FlgN, FliS and FliT in complex with their cognate filament-type substrates (Extended Data Fig. 2)^7–11^. These two domains are connected by a flexible linker (FlhA_L_) that has been implicated in both export gate activation and substrate specificity switching^12,13^ (Fig. 1a). Genetic and biochemical studies suggest that the C-terminal region of FlhA_L_ regulates conformational changes of the FlhA ring^14,15^, but the structural basis of this regulation remains unknown.

**Fig. 1.**
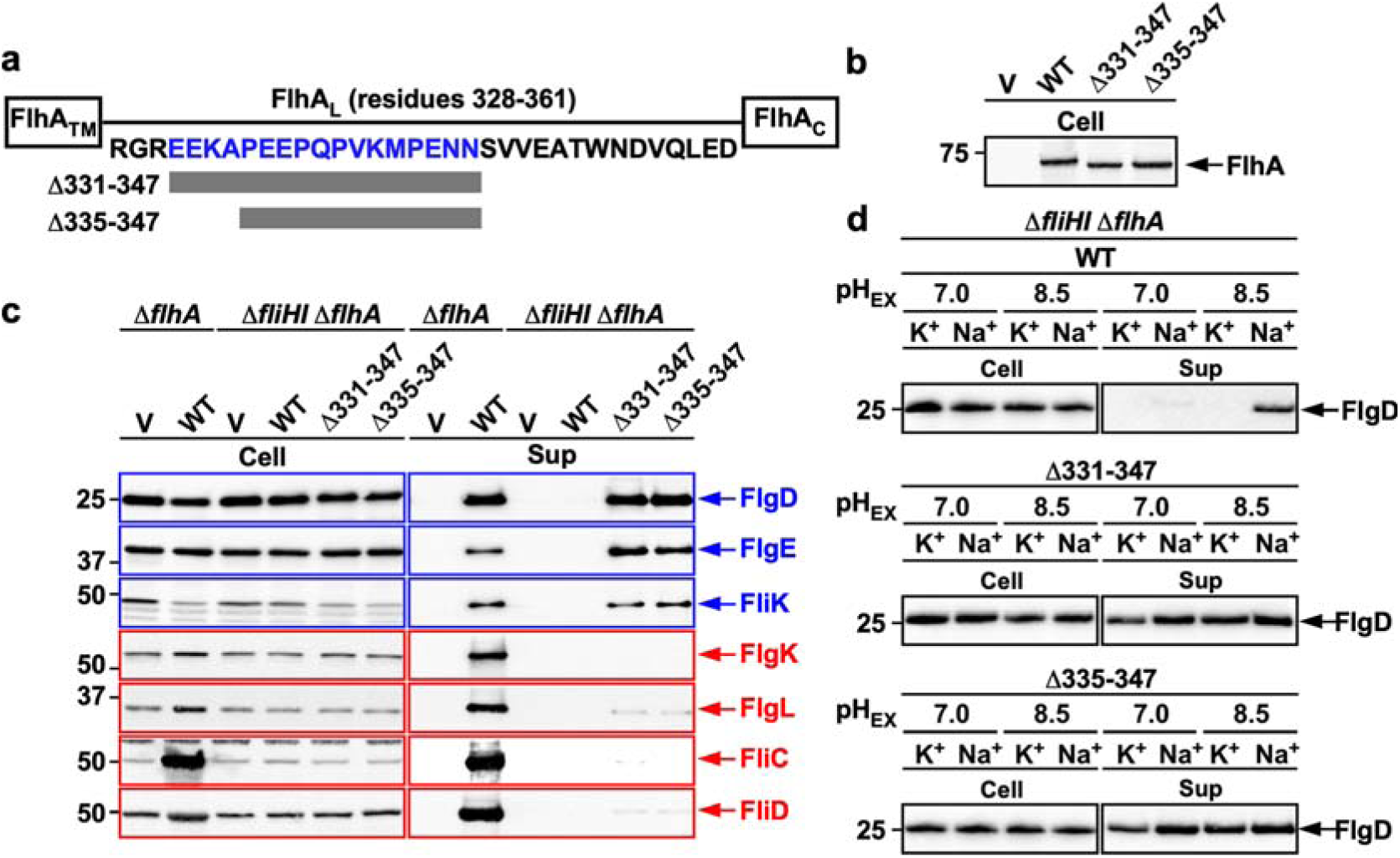
FlhA linker deletions lock the export gate in an autonomously activated hook-type state. **(a)** Primary sequence of the *Salmonella* FlhA linker region (FlhA_L_) connecting the N-terminal transmembrane domain (FlhA_TM_) and the cytoplasmic domain (FlhA_C_) (UniProt: P40729). FlhA_L_ is predicted to comprise residues 328–361. **(b)** Steady-state expression levels of FlhA in *flhA* linker deletion mutants. Whole-cell (Cell) fractions were prepared from *Salmonella* NH003 (Δ*fliH-fliI* Δ*flhA*) carrying pTrc99AFF4 (V), pMM130 (WT), pMKM130-207 (Δ331–347) or pMKM130-212 (Δ335–347). Samples were normalized to OD_600_ and analysed by SDS-PAGE and immunoblotting with a polyclonal anti-FlhA_C_ antibody. **(c)** Secretion profiles of *flhA* linker deletion mutants (Δ331–347 and Δ335–347) under ATPase-deficient conditions. Whole-cell (Cell) and culture supernatant (Sup) fractions were prepared from *Salmonella* NH001 (Δ*flhA*) carrying pTrc99AFF4 (V) or pMM130 (WT), and NH003 (Δ*fliHI* Δ*flhA*) carrying pTrc99AFF4 (V), pMM130 (WT), pMKM130-207 (Δ331–347) or pMKM130-212 (Δ335–347), and analysed by immunoblotting with the indicated polyclonal antibodies. Hook-type and filament-type substrates are highlighted in blue and red, respectively. Molecular mass markers (kDa) are indicated. Three independent experiments were performed. **(d)** Na^+^- and membrane potential-dependent FlgD secretion in the absence of FliH and FliI. Immunoblots of whole-cell (Cell) and culture supernatant fractions from *Salmonella* NH003 carrying pMM130 (top), pMKM130-207 (middle) or pMKM130-212 (bottom), grown at 30□°C in T-broth at pH 7.0 or 8.5 with either 100□mM K^+^ or Na^+^. Molecular mass markers (kDa) are indicated. Three independent experiments were performed.

Despite recent advances in cryo-electron microscopy (cryo-EM), the mechanism by which FlhA couples ion-driven export to assembly stage-dependent substrate selection remains unclear. Structures of the cytoplasmic FlhA_C_ ring and the FliPQR-FlhB protein-export channel complex have been determined at near-atomic resolution^16–19^; however, FlhA_TM_ has remained unresolved owing to instability and conformational heterogeneity of its ring. Consequently, how the cytoplasmic substrate-docking platform is functionally linked to the membrane-embedded export engine—and how this linkage is coordinated with substrate specificity switching—remains a major unanswered question.

Here we identify FlhA linker mutations that autonomously activate the export gate and lock the system in a functionally active, hook-type substrate export state. Using these mutants, we reconstitute and determine a high-resolution structure of the full-length FlhA ring complex by cryo-EM. The structure shows that FlhA_L_-mediated docking of the FlhA_C_ ring to FlhA_TM_ stabilizes the FlhA_TM_ ring as the export engine while sterically excluding filament-type substrate binding to FlhA_C_. These findings reveal a structural mechanism by which the membrane-embedded export engine is allosterically coupled to stage-specific substrate selection during flagellar assembly.

## Results

### Linker deletions lock FlhA in an autonomously active hook-type state

In situ structural analysis by cryo-electron tomography has shown that the FlhA_C_ ring projects into the central cavity of the basal body C-ring, positioning it distal from the transmembrane export gate^9,20^. Because the export gate can be autonomously activated by an increase in membrane potential even in the absence of FliH and FliI^4^, we hypothesized that FlhA_C_ associates more closely with FlhA_TM_ through a linker-mediated conformational change in the activated state.

To test this hypothesis, we constructed a series of FlhA linker deletion mutants and analyzed their secretion profiles under ATPase-deficient conditions. For this purpose, we used two *Salmonella* strains, Δ*fliH-fliI* Δ*flhA* and Δ*fliH-fliI flhB*(P28T) Δ*flhA*, as hosts. The *flhB*(P28T) mutation is known to increase the probability of flagellar assembly in the absence of FliH and FliI^21^. Secretion assays were performed by immunoblotting. Among the mutants tested, two linker deletion variants (Δ331–347 and Δ335–347) showed no changes in the steady cellular level of FlhA (Fig. 1b) but exhibited markedly elevated export activities (Fig. 1c and Extended Data Fig. 3a). Notably, these mutants efficiently secreted hook-type substrates (FlgD, FlgE and FliK) but failed to secrete filament-type substrates (FlgK, FlgL, FliC and FliD) (Fig. 1c), indicating a defect in substrate specificity switching. Consistently, pull-down assays using GST affinity chromatography revealed that the FlhA linker deletions inhibit the binding of the filament-type export chaperone FlgN to FlhA (Extended Data Fig. 4).

To investigate the mechanism underlying this autonomous activation, we examined the dependence of protein export on membrane potential under ATPase-deficient conditions (Fig. 1d and Extended Data Fig. 3b). Because the membrane potential increases with external pH^4^, FlgD secretion was measured at pH 7.0 and 8.5 in the presence of either 100 mM KCl or 100 mM NaCl. As expected, the Δ*fliH-fliI* strain showed no detectable export activity of FlgD at pH 7.0 in the absence of Na^+^, regardless of the presence of the *flhB*(P28T) mutation. At pH 8.5, the Δ*fliH-fliI flhB*(P28T) strain exhibited a nearly full export activity in the presence of Na^+^ but only a low export activity in the absence of Na^+^, and the Δ*fliH-fliI* strain did not show any activity, consistent with previous observations^4^. In contrast, both linker deletion mutants showed a robust export activity even at pH 7.0, and this activity was further enhanced by Na⁺ in a manner similar to wild-type FlhA. These results indicate that the linker deletions stabilize a functionally active state of the dual-fuel export engine.

Previous studies have shown that cooperative remodeling of the FlhA_C_ ring during substrate specificity switching depends on FliH and FliI^22,23^. We therefore examined the phenotypes of the linker deletion mutants in the presence of FliH and FliI. Both mutants were non-motile on soft agar plates (Extended Data Fig. 5a). They secreted substantially higher levels of hook-type substrates than wild-type cells (Extended Data Fig. 5b) and produced polyhooks (Extended Data Fig. 5c), but failed to secrete filament-type substrates (Extended Data Fig. 5b), resulting in the absence of filaments (Extended Data Fig. 5d).

These results demonstrate that the linker deletions lock FlhA in a functionally active hook-type export state that is competent for hook-type substrate export but defective in switching to filament-type export.

### In vitro reconstitution of FlhA ring complex in peptidisc

To enable efficient purification of full-length FlhA, we introduced an N-terminal His-tag, which has been shown to have no detectable effect on FlhA function^24^. His-tagged wild-type FlhA and the linker deletion variant (Δ331–347) were overproduced in a *Salmonella* strain lacking the *flhDC* master operon (SJW1368), and membrane fractions were solubilized with lauryl maltose neopentyl glycol (LMNG). The proteins were purified by Ni-affinity chromatography, followed by size exclusion chromatography.

Under these conditions, wild-type FlhA did not form ring-like particles in the presence of LMNG over a wide range of pH (Extended Data Fig. 6), suggesting that detergent solubilization does not support stable assembly of the full-length complex. To overcome this limitation, we replaced LMNG with peptidisc composed of short amphipathic peptides that wrap around the transmembrane region and shield hydrophobic surfaces^25^. We then assessed FlhA ring formation across a range of pH conditions using negative-stain electron microscopy (Fig. 2a and Extended Data Fig. 6).

**Fig. 2.**
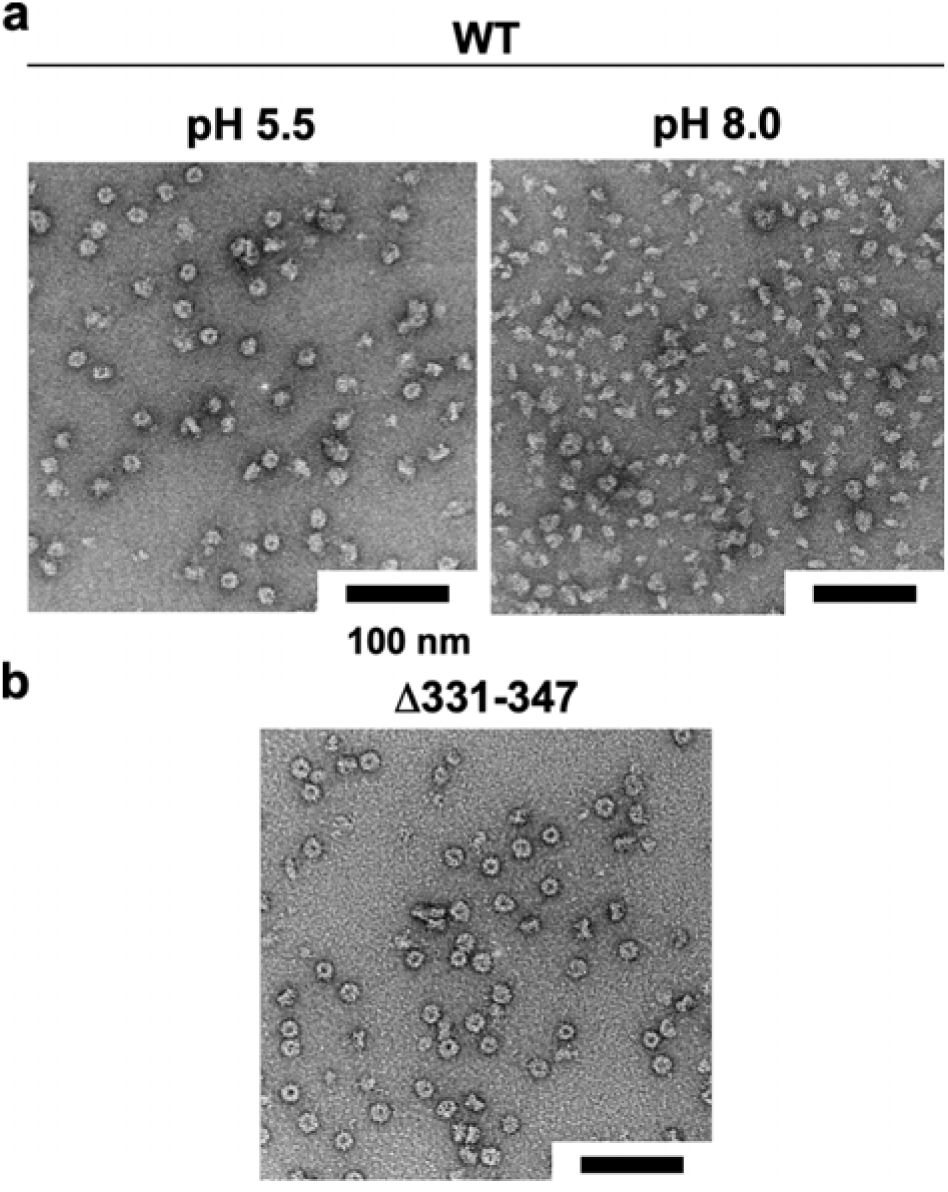
Effect of pH on FlhA ring formation in peptidisc. **(a)** Reconstitution of the wild-type FlhA ring in peptidisc. Purified wild-type FlhA solubilized in LMNG was concentrated to 2.5□mg□ml⁻¹ and buffer-exchanged using Amicon Ultra centrifugal filters. The following buffers were used: 50□mM piperazine-HCl (pH 5.5) and 50□mM Tris-HCl (pH 8.0), each containing 150□mM NaCl and 0.005%(w/v) LMNG. LMNG was then replaced with peptidisc under each pH condition. Scale bar, 100□nm. **(b)** Reconstitution of the FlhA(Δ331–347) ring in peptidisc at pH 5.5.

Wild-type FlhA efficiently formed ring-like particles at pH 5.5, close to its isoelectric point, whereas the frequency of ring formation decreased with increasing pH. At lower pH, peptidisc-reconstituted FlhA showed increased aggregation. The linker deletion mutant FlhA(Δ331–347) also efficiently formed ring-like particles under the same pH 5.5 condition (Fig. 2b).

These results indicate that peptidisc reconstitution enables assembly of the full-length FlhA ring complex and suggest that electrostatic intersubunit interactions influence its structural stability. Together, these data establish conditions for structural analysis of the full-length FlhA ring complex.

### Cryo-EM structures reveal state-dependent coupling between FlhA_C_ and FlhA_TM_

To investigate how the linker deletion locks the FlhA ring in a functionally active hook-type state, we performed high-resolution cryo-EM analysis of the wild-type FlhA ring and its linker deletion mutant (Δ331–347) (Extended Data Figs 7 and 8 and Extended Data Table 1). Single-particle analysis of both samples yielded clear 2D class averages displaying well-defined ring structures consistent with nonameric ring assembly (Fig. 3 and Extended Data Fig. 9).

**Fig. 3.**
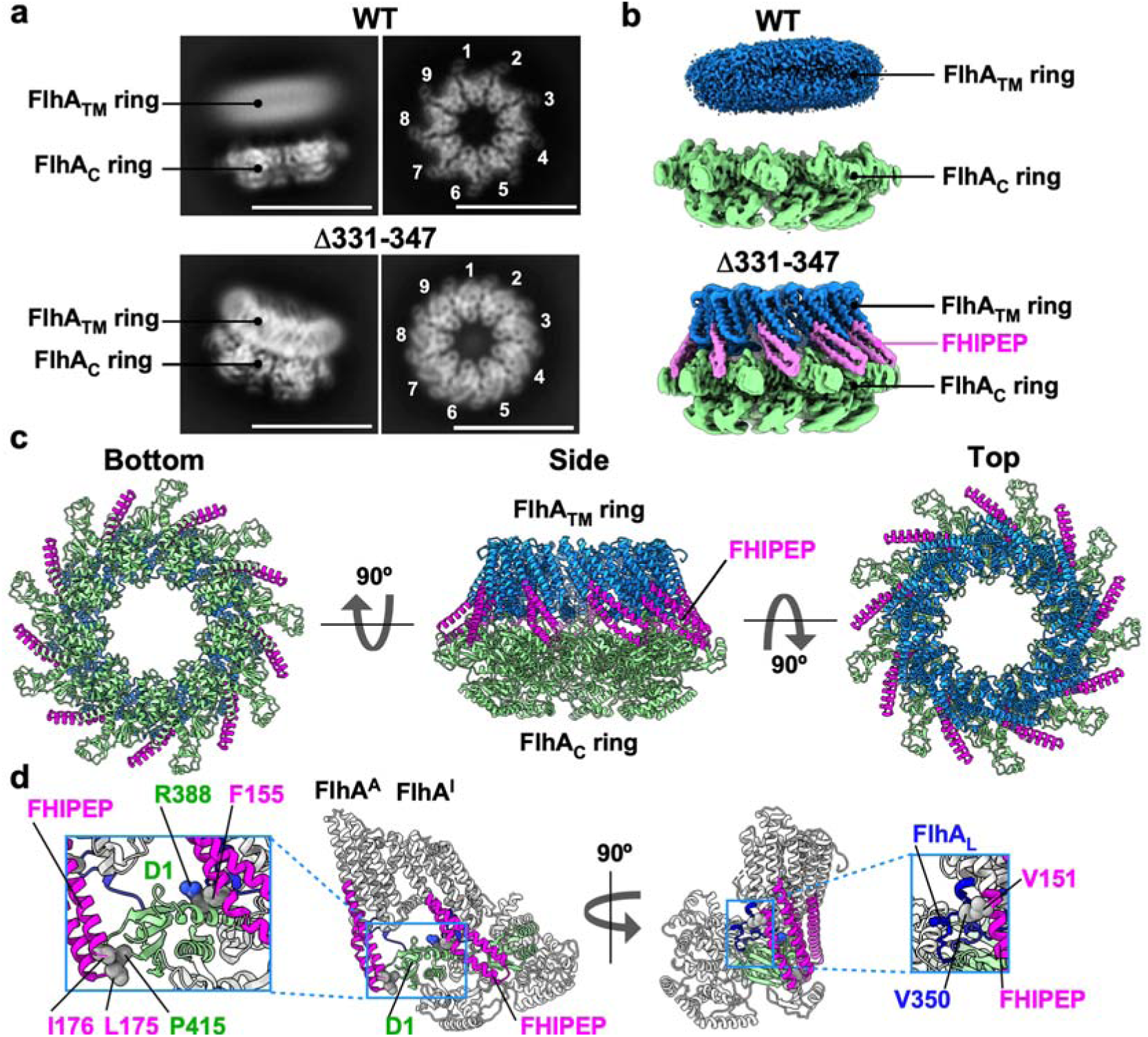
Cryo-EM structure reveals FlhA_L_-mediated docking of the FlhA_C_ ring onto the transmembrane ring. **(a)** Representative 2D class averages of wild-type (WT) and FlhA linker deletion mutant (Δ331–347) ring complexes obtained by single-particle cryo-EM image analysis. FlhA assembles into a homononameric ring. Scale bar, 15 nm. **(b)** Cryo-EM density maps of WT and FlhA(Δ331–347) ring complexes. In the WT complex, the cytoplasmic FlhA_C_ ring is well resolved, whereas the transmembrane FlhA_TM_ ring is poorly defined, indicating substantial conformational heterogeneity. In contrast, the linker mutant shows well-resolved densities for both the FlhA_C_ and the FlhA_TM_ rings. **(c)** Cα ribbon representation of the atomic model of the FlhA(Δ331–347) ring complex (PDB ID: 25ZD). The FlhA_TM_, FHIPEP (residues 141–208), and FlhA_C_ domains are colored dodger blue, magenta, and light green, respectively. The FHIPEP region bridges FlhA_TM_ and FlhA_C_. **(d)** Close-up view of the interaction of the FHIPEP region (magenta) with FlhA_L_ (blue) and domain D1 of FlhA_C_ (light green).

In the wild-type structure, FlhA_L_ adopts an extended conformation that spatially separates the cytoplasmic FlhA_C_ ring from the transmembrane FlhA_TM_ ring (Fig. 3a), consistent with previous in situ observations^9,20^. Following iterative 3D refinement with C1 symmetry, a 3D reconstruction of the wild-type FlhA ring was obtained. While the FlhA_C_ ring was well resolved, the transmembrane region remained poorly defined (Fig. 3a, upper left), even after chemical crosslinking with glutaraldehyde (Extended Data Fig. 10), indicating substantial conformational heterogeneity. In other words, the FlhA_TM_ ring appears to adopt multiple conformational states. In addition, weak but detectable density possibly corresponding to FlhA_L_ was observed in the wild-type FlhA ring reconstructed with C9 symmetry (Extended Data Fig. 11), supporting the notion that FlhA_L_ dynamically connects the FlhA_TM_ and FlhA_C_ rings even in the resting state.

In contrast, in the linker deletion mutant, the FlhA_C_ ring is docked onto the well-ordered FlhA_TM_ ring. Accordingly, clear helical features corresponding to transmembrane helices of FlhA_TM_ were readily visible in the 2D class averages, unlike in the wild-type (Fig. 3a, lower left). Subsequent 3D refinement with C9 symmetry yielded a reconstruction of the FlhA(Δ331–347) ring at 2.73 Å resolution (EMD-80488) (Fig. 3b, lower panel and Extended Data Fig. 8c). The map reveals that docking of the cytoplasmic ring stabilizes the transmembrane helices within the FlhA_TM_ ring, resulting in a well-ordered conformation. The improved resolution further enabled accurate atomic model building of the full-length complex (PDB ID: 25ZD) (Fig. 3c).

Residues 141–208 of FlhA form a highly conserved small cytoplasmic domain termed FHIPEP (flagellum/<u>h</u>ypersensitive response/invasion <u>p</u>rotein <u>e</u>xport <u>p</u>ore)^26^. This domain has been implicated in both H⁺-coupled protein export^15,27^ and substrate specificity switching from hook-type to filament-type^28^. In our structure, the FHIPEP domain is located in a gap formed between adjacent FlhA_C_ subunits and makes hydrophobic contacts with both FlhA_L_ and domain D1 of FlhA_C_, thereby mediating docking of the FlhA_C_ ring to the FlhA_TM_ ring (Fig. 3d). Three major hydrophobic contact sites were identified. The conserved Leu175 and Ile176 residues interact intramolecularly with Pro415 in domain D1, whereas the conserved Phe155 residue forms an intermolecular contact with Arg388 in domain D1. In addition, the conserved Val151 residue interacts with Val350 in FlhA_L_. These residues have been shown to be functionally important, and Val151 has been implicated in substrate specificity switching^28^. Given that the linker deletion mutant is defective in switching to filament-type export, these observations suggest that FHIPEP-mediated hydrophobic interactions stabilize the hook-type state of the FlhA ring.

Together, these observations indicate that FlhA_L_-mediated docking of the FlhA_C_ ring stabilizes the FlhA_TM_ ring structure and underlies a state-dependent coupling between the cytoplasmic and transmembrane regions of FlhA.

### Structure of the FlhA transmembrane domain

We built an atomic model of the FlhA_TM_ ring with C9 symmetry. However, the TM1 and TM7 helices were not resolved (Fig. 4a, left and Extended Data Table 2), indicating conformational heterogeneity in these regions. To address this, we performed 3D classification and refinement with C1 symmetry (Extended Data Fig. 8b), which yielded two distinct maps at 3.39 Å (Class 1, EMD-80489) and 3.43 Å resolution (Class 2, EMD-80490) (Extended Data Fig. 8d and Extended Data Table 1). These maps enabled us to build atomic models of TM1 and TM7 (Class 1, PDB ID: 25ZG; Class 2, PDB ID: 25ZI) (Fig. 4a, right and Extended data Table 2).

**Fig. 4.**
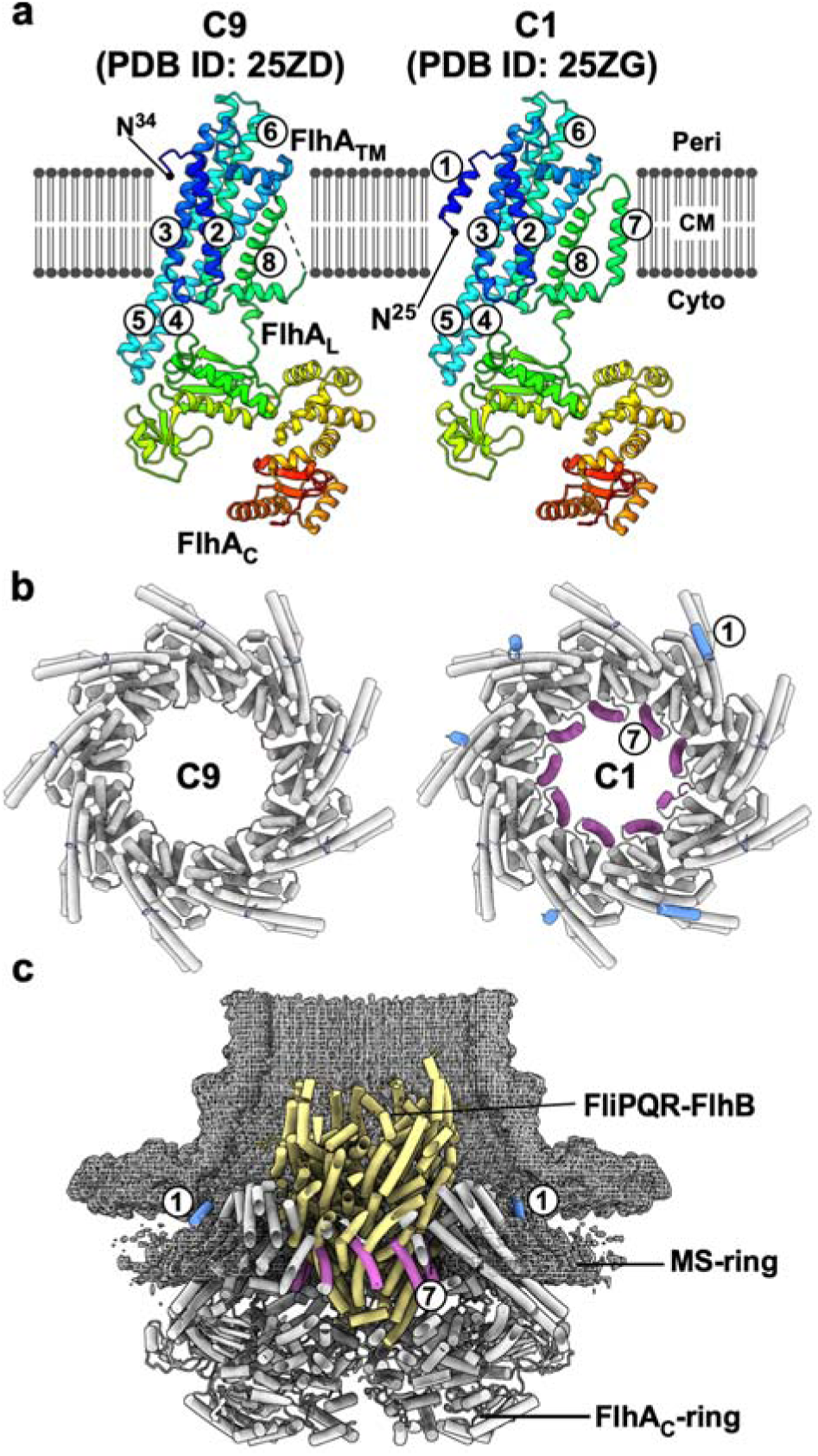
Cryo-EM structure of the FlhA transmembrane ring. **(a)** Membrane topology of the FlhA subunit within the ring. The transmembrane domain of FlhA contains ten α-helices (see Extended Data Table 2). Helices α1, α2, α3, α5, α6, α7, α9 and α10 correspond to transmembrane helices TM1–TM8, respectively. TM1 and TM7 are not resolved in the C9-symmetrized structure (PDB ID: 25ZD). In contrast, TM1 is visible in subunits A, D, E, G and H, but not in subunits B, C, F and I, in the C1 reconstruction (PDB ID: 25ZG). TM7 is resolved in all FlhA subunits. The α5 and α6 helices also extend into the cytoplasm to form a cytoplasmic FHIPEP domain. **(b)** Location of TM1 and TM7 within the FlhA_TM_ ring. TM7 (orchid) is positioned at the innermost region of the ring, whereas TM1 (cornflower blue) is located at the outermost region. **(c)** Model of the FlhA_TM_ ring within the MS-ring. TM1 is proposed to interact with the MS-ring (EMD-60007) and stabilize the export gate complex during rapid protein export. TM7 is positioned adjacent to the FliPQR–FlhB complex (khaki, PDB ID: 6S3L) and may couple inward ion flow to outward protein translocation through the FliPQR export channel.

In these structures, TM7 is positioned at the innermost region of the FlhA_TM_ ring, whereas TM1 is located at the outermost region (Fig. 4b). Given that the FliPQR-FlhB complex occupies the central pore of the FlhA_TM_ ring (Fig. 4c), TM7 is likely to interact with this complex and adopt a stabilized conformation. Because the FlhA_TM_ ring functions as a transmembrane ion channel capable of conducting both H^+^ and Na^+5^, TM7 may couple inward ion flow through the FlhA channel to outward protein translocation through the FliPQR export channel. Conversely, TM1 is positioned at the periphery of the ring and may interact with the MS-ring (Fig. 4c). TM1 therefore likely contributes to stabilization of the export gate complex during rapid protein export. Together, these structures reveal a well ordered conformation of the ion-conducting export engine.

### FHEIPEP-mediated docking blocks filament-type substrate binding

Flagellar export chaperones in complex with their cognate filament-type substrates bind to a well-conserved dimple located at the interface between domains D1 and D2 of FlhA_C_^8,10,11^. The conserved residues Asp456, Phe459 and Thr490 are directly involved in this chaperone-FlhA_C_ interaction.

To examine structural differences between hook-type and filament-type states, we performed 3D refinement of the wild-type FlhA_C_ ring with either C1 or C9 symmetry (Extended Data Fig. 7b), yielding atomic models at 3.3 Å (EMD-80484, PDB ID: 25YY) and 3.05 Å resolution (EMD-80485, PDB ID: 25YZ), respectively (Extended Data Fig. 7c and Extended Data Table 3). These resolutions were lower than that of the FlhA(Δ331–347) ring, suggesting conformational variability among FlhA_C_ subunits. Therefore, the FHIPEP-FlhA_C_ interactions seem likely to stabilize the FlhA_C_ ring conformation.

To reduce this heterogeneity, we performed 3D refinement of chemically crosslinked wild-type FlhA_C_ rings, which yielded improved maps at 2.87 Å resolution in C1 symmetry (EMD-80486, PDB ID: 25ZA) and 2.6 Å resolution in C9 symmetry (EMD-80487, PDB ID: 25ZB) (Extended Data Figs. 10 and 12a and Extended Data Table 3). The overall structure of each FlhA_C_ subunit was highly similar to that observed in the FlhA(Δ331–347) ring (Extended Data Fig. 12b), with both adopting an open conformation comparable to that of the previously reported 3A5I structure.

To understand why the FlhA(Δ331–347) mutant fails to produce filaments, we superimposed the crystal structure of the FlhA_C_-FliS-FliC chaperone-substrate complex onto the FlhA(Δ331–347) ring. This analysis revealed that the FHIPEP domain sterically blocks docking of the FliS-FliC complex onto FlhA_C_ (Fig. 5a, left). Consistent with the structural model, pull-down assays using GST affinity chromatography revealed that this FlhA linker deletion inhibits binding of the filament-type FlgN export chaperone to FlhA (Extended Data Fig. 4). On the other hand, the FliS-FliC complex can be accommodated within the wild-type FlhA_C_ ring (Fig. 5a, right). These observations provide a clear structural basis for hook-type substrate specificity.

**Fig. 5.**
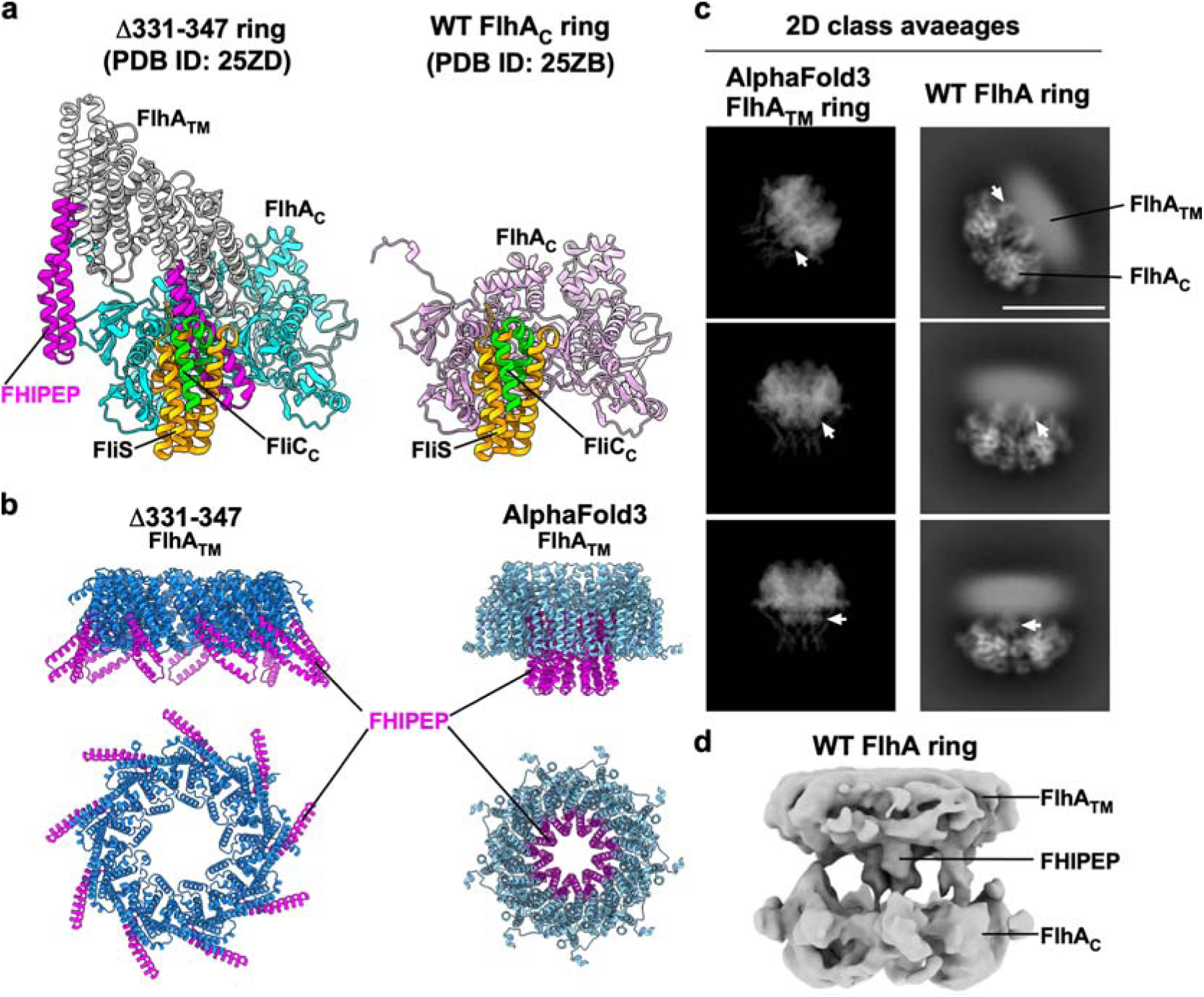
FHIPEP inhibits docking of the FliS-FliC chaperone-substrate complex onto the FlhA_C_ ring. **(a)** Superposition of the crystal structure of the FlhA_C_-FliS-FliC_C_ complex [PDB ID: 6CH3; FliS, orange; C-terminal region of FliC (FliC_C_), lime] onto the FlhA(Δ331–347) ring (left panel) [PDB ID: 25ZD; FlhA_TM_, light grey; FHIPEP, magenta; FlhA_C_, cyan] and the chemically crosslinked wild-type FlhA_C_ ring (right panel) [PDB ID: 25ZB; FlhA_C_, plum]. In the FlhA(Δ331–347) ring, the FHIPEP domain sterically blocks docking of the FliS–FliC complex onto FlhA_C_. **(b)** Structural comparison between the transmembrane FlhA(Δ331–347) ring and the AlphaFold3-predicted FlhA transmembrane ring. The predicted model exhibits a distinct transmembrane topology in which the FHIPEP domain is repositioned towards the interior of the ring. **(c)** Comparison between experimentally obtained 2D class averages of the wild-type FlhA ring and 2D projections generated from the AlphaFold3-predicted FlhA_TM_ ring model. White arrows indicate FHIPEP-like density extending between the FlhA_TM_ and FlhA_C_ rings. Scale bar, 15 nm. **(d)** Cryo-EM density map of the wild-type FlhA ring reconstructed from 19,640 particles by single-particle cryo-EM analysis. Density extending from the center of the FlhA_TM_ ring into the central pore of the FlhA_C_ ring is consistent with repositioning of the FHIPEP domain.

Because the FHIPEP domain is positioned on the outer surface of the FlhA ring in the hook-type state, it must be repositioned during the transition to the filament-type state. Consistent with this idea, an AlphaFold3-predicted model of the FlhA_TM_ ring exhibits a distinct topology in which the FHIPEP domain is relocated to the interior of the ring (Fig. 5b).

To further investigate topological remodeling of the FlhA_TM_ ring, we selected 2D class averages of the wild-type FlhA that exhibited prominent FHIPEP-like structural features extending from the center of the FlhA_TM_ ring (Extended Data Fig. 13a). Comparison of 2D projections generated from the predicted model with experimentally obtained 2D class averages suggests that a FHIPEP-like density extends into the cytoplasmic space between the FlhA_TM_ and FlhA_C_ rings (Fig. 5c).

Furthermore, refinement of the wild-type FlhA ring with C9 symmetry using 19,640 particles yielded a 3D cryo-EM map at 8.34 Å resolution (Extended Data Fig. 13b). This map revealed densities extending towards the central cavity of the FlhA_C_ ring (Fig. 5d). Given the conformational heterogeneity of the FlhA_TM_ ring in the resting state, these observations suggest that structural flexibility enables repositioning of the FHIPEP domain during the hook-to-filament transition. Such rearrangement would relieve steric hindrance, thereby allowing filament-type substrate binding and initiation of filament assembly.

This model is further supported by genetic evidence showing that the FHIPEP domain is directly involved in substrate specificity switching and that mutations within the FHIPEP region induce topological rearrangements of the FlhA_TM_ ring (Extended Data Fig. 14)^28^.

## Discussion

In this study, we determined the structure of the FlhA ring in a functionally active, hook-type export state and resolved the transmembrane domain and its nonameric ring that has remained the final missing structural information of the fT3SS. By capturing this structure using linker mutations that autonomously activate the export gate, we provide a structural view of the full-length FlhA ring in an active conformation.

Our structure reveals that FlhA_L_-mediated docking of the FlhA_C_ ring onto the FlhA transmembrane domain stabilizes the FlhA_TM_ ring as the ion-driven export engine of the fT3SS. In the wild-type complex, the FlhA_TM_ ring appears structurally heterogeneous, whereas in the linker mutant it adopts a well-ordered conformation (Fig. 3a,b). These observations suggest that docking of the cytoplasmic ring promotes structural ordering of the transmembrane helices of FlhA, thereby stabilizing the FlhA_TM_ ring to activate the export engine. Thus, direct coupling between the cytoplasmic and transmembrane domains emerges as a key feature of the FlhA function.

A central finding of this study is that the structural rearrangement that stabilizes the export engine also controls substrate specificity. The FHIPEP region mediates docking of the FlhA_C_ ring onto the transmembrane domain (Fig. 3d), while simultaneously occluding binding of the FliS-FliC chaperone-substrate complex sterically (Fig. 5a). This dual role provides a direct structural explanation as to why the captured state supports efficient export of hook-type substrates while preventing transition to filament-type export. These results indicate that export activation and substrate specificity switching are intrinsically coupled through a single conformational transition.

Our findings establish FlhA_L_ as an allosteric coupler that integrates ion-driven export activation with assembly-stage-dependent substrate selection. In this model, linker-mediated remodeling of the FlhA ring defines discrete functional states of the fT3SS, coordinating energy transduction with ordered protein export during flagellar assembly. More broadly, this work provides a structural framework for understanding how the membrane-embedded export engines couple their conformational dynamics to substrate selection. Given the conservation of type III secretion systems, similar coupling principles must be operating in virulence-associated injectisomes and related protein export systems.

## Acknowledgements

We acknowledge Fumiaki Makino, Tomoko Miyata, Yasuyo Abe, and Yoshie Kushima for technical assistance and Katsumi Imada for continuous support and encouragement. This work was supported in part by JSPS KAKENHI Grant Numbers JP22K06162 and JP26K09266 (to M.K.), JP22H02573 and JP26K01964 (to T.M.) and MEXT KAKENHI Grant Number JP26H01683 (to T.M.). This work has also been supported by Research Support Project for Life Science and Drug Discovery (BINDS) from AMED under Grant Number JP23am121003, JP24am121003, and JP25am121003 (to K.N.), by the Cyclic Innovation for Clinical Empowerment (CiCLE) from AMED under Grant Number JP17pc0101020 (to K.N.), and by JEOL YOKOGUSHI Research Alliance Laboratories of The University of Osaka (to K.N.).

## Author Contributions

T.M. and K.N. conceived and designed research; M.K. and H.T. constructed *flhA* linker deletion mutants; M.K. prepared samples for cryo-EM and collected cryo-EM image data; M.K. and K.K. analysed cryo-EM image data; M.K. built atomic models; T.M. performed genetic, biochemical, and physiological experiments; T.M. and K.N. wrote the paper based on discussion with other authors.

## Competing interests

The authors declare no competing interests.

## Data availability

The cryo-EM density maps have been deposited in the Electron Microscopy Data Bank (EMDB) under accession codes EMD-80484 for the wild-type FlhA_C_ ring reconstructed with C1 symmetry, EMD-80485 for the wild-type FlhA_C_ ring reconstructed with C9 symmetry, EMD-80486 for the chemically crosslinked wild-type FlhA_C_ ring reconstructed with C1 symmetry, EMD-80487 for the chemically crosslinked wild-type FlhA_C_ ring reconstructed with C9 symmetry, EMD-80488 for the FlhA(Δ331–347) ring reconstructed with C9 symmetry, and EMD-80489 (Class 1) and EMD-80490 (Class 2) for the FlhA(Δ331–347) ring reconstructed with C1 symmetry.

Atomic coordinates have been deposited in the Protein Data Bank (PDB) under accession codes 25YY and 25YZ for the wild-type FlhA_C_ ring reconstructed with C1 and C9 symmetry, respectively; 25ZA and 25ZB for the chemically crosslinked wild-type FlhA_C_ ring reconstructed with C1 and C9 symmetry, respectively; 25ZD for the FlhA(Δ331–347) ring reconstructed with C9 symmetry; and 25ZG and 25ZI for Class 1 and Class 2 of the FlhA(Δ331–347) ring reconstructed with C1 symmetry, respectively.

All data supporting the findings of this study are available within the paper and its Supplementary Information. Strains, plasmids, polyclonal antibodies and other materials are available from the corresponding author upon reasonable request.

## METHODS

### *Salmonella* strains, plasmids, DNA manipulations, and media

*Salmonella* strains and plasmids used in this study are listed in Extended Data Table 4. DNA manipulations were performed using standard protocols. The cloned DNA fragments were confirmed by DNA sequencing (Eurofins Genomics). L-broth contained 1% (w/v) tryptone, 0.5% (w/v) yeast extract, and 0.5% (w/v) NaCl. T-broth contained 1% (w/v) Bacto tryptone, 10 mM potassium phosphate. The pH of T-broth was adjusted to the desired final pH by addition of KOH. 2×YT medium contained 1.6% (w/v) tryptone, 1.0% (w/v) yeast extract, 0.5% (w/v) NaCl. Soft tryptone agar plates contained 1% (w/v) tryptone, 0.5% (w/v) NaCl, and 0.35% (w/v) agar. Ampicillin was added as needed at a final concentration of 100 μg ml^-1^.

### Secretion assays

*Salmonella* cells were grown in L-broth supplemented with ampicillin at 30°C with shaking until the optical density at 600 nm (OD_600_) reached approximately 1.2–1.4. For analysis of Na⁺- and membrane potential-dependent protein secretion, 50 μl of the overnight culture was inoculated into 5 ml of fresh T-broth adjusted to pH 7.0 or 8.5 and supplemented with either 100 mM KCl or 100 mM NaCl and grown at 30°C with shaking until the OD_600_ reached approximately 1.4–1.6.

Cells were then harvested by centrifugation to separate cell pellets and culture supernatants. Protein samples from whole-cell and culture supernatant fractions were normalized based on the OD_600_ of each culture to ensure equivalent cell numbers. Cell pellets were directly resuspended in SDS loading buffer [62.5 mM Tris-HCl (pH 6.8), 2% (w/v) sodium dodecyl sulfate (SDS), 10% (w/v) glycerol, and 0.001% (w/v) bromophenol blue] supplemented with 1 μl of 2-mercaptoethanol. Proteins in the culture supernatants were precipitated with 10% (v/v) trichloroacetic acid, resuspended in Tris–SDS loading buffer (one volume of 1 M Tris mixed with nine volumes of SDS loading buffer) containing 1 μl of 2-mercaptoethanol, and heated at 95°C for 3 min.

Protein samples were separated by SDS-polyacrylamide gel electrophoresis (SDS-PAGE) and transferred to nitrocellulose membranes (Cytiva) using a transblotting apparatus (Hoefer). Then, immunoblotting with polyclonal anti-FlgD, anti-FlgE, anti-FliK, anti-FlgK, anti-FlgL, anti-FliC, anti-FliD, or anti-FlhA_C_ antibody as the primary antibody and anti-rabbit IgG, HRP-linked whole Ab Donkey (Cytiva) as the secondary antibody was carried out. Immunoblotting was performed with an iBind Flex Western Device according to the manufacturer’s instructions (Thermo Fisher Scientific). Chemiluminescent signals were detected using Amersham ECL Prime Western Blotting Detection Reagent (Cytiva) and captured with a Luminoimage Analyzer LAS-3000 (GE Healthcare). Image data were processed using Photoshop software (Adobe). The assay was performed at least three times to confirm the reproducibility of the results.

### Pull-down assays by GST affinity chromatography

Cell lysates prepared from SJW1368 (Δ*cheW-flhD*) expressing GST-FlgN were mixed with lysates prepared from NH003 (Δ*fliH-fliI flhA*) transformed with pMM130 (wild-type FlhA), pMKM130-207 [FlhA(Δ331–347)], or pMKM130-212 [FlhA(Δ335–347)]. The mixtures were loaded onto a glutathione Sepharose 4B column (bed volume, 1 ml; GE Healthcare). After washing the column with 5 ml PBS (8 g of NaCl, 0.2 g of KCl, 3.63 g of Na_2_HPO_4_•12H_2_O, 0.24 g of KH_2_PO_4_, pH 7.4 per liter), bound proteins were eluted with 5 ml buffer containing 50 mM Tris-HCl (pH 8.0) and 10 mM reduced glutathione. Fractions containing GST-FlgN were analyzed by SDS-PAGE followed by Coomassie Brilliant Blue (CBB) staining and immunoblotting using a polyclonal anti-FlhA_C_ antibody. At least three independent pull-down assays were performed.

### Hook-length measurements

*Salmonella* cells were grown in 500 ml of L-broth containing ampicillin at 30°C with shaking until the cell density had reached an OD_600_ of ca. 1.0. After centrifugation (10,000 g, 10 min, 4°C), the cells were suspended in 20 ml of ice-cold sucrose buffer containing 0.1 M Tris-HCl (pH 8.0) and 0.5 M sucrose. EDTA and lysozyme were added at the final concentrations of 10 mM and 0.1 mg ml^-1^, respectively. The cell suspensions were stirred for 30 min at 4°C, and Triton X-100 and MgSO_4_ were added at final concentrations of 1% (w/v) and 10 mM, respectively. After stirring on ice for 1 h, cell lysates were adjusted to pH 10.5 with 5 M NaOH and then centrifuged (10,000 g, 20 min, 4°C) to remove cell debris. After ultracentrifugation (45,000 g, 60 min, 4°C), pellets were resuspended in buffer A containing 10 mM Tris-HCl (pH 8.0), 5 mM EDTA, and 1% (w/v) Triton X-100, and this solution was loaded onto a 20–50% (w/w) sucrose density gradient in buffer A. After ultracentrifugation (49,100 g, 13 h, 4°C), hook-basal bodies and polyhook-basal bodies with or without filaments were collected and ultracentrifuged (60,000 g, 60 min, 4°C). Pellets were suspended in acidic buffer containing 50 mM glycine (pH 2.5) and 0.1% (w/v) Triton X-100 to depolymerize flagellar filaments. After ultracentrifugation (60,000 g, 60 min, 4°C), pellets were resuspended in 50 μl of buffer A.

Samples were negatively stained with 2% (w/v) uranyl acetate. Electron micrographs were taken using JEM-1400Flash (JEOL, Tokyo, Japan) operated at 100 kV. The length of hooks and polyhooks was measured by ImageJ version 1.54p (National Institutes of Health).

### Fluorescence microscopy

*Salmonella* cells were grown in 5 ml of L-broth containing ampicillin. The cells were attached to a cover slip (Matsunami glass, Japan), and unattached cells were washed away with motility buffer containing 10 mM potassium phosphate (pH 7.0), 0.1 mM EDTA, and 10 mM sodium □-lactate. Then, flagellar filaments were labelled using anti-FliC antibody and anti-rabbit IgG conjugated with Alexa Fluor 594 (Invitrogen). After washing twice with motility buffer, the cells were observed by an inverted fluorescence microscope (IX-83, Olympus) with a 150× oil immersion objective lens (UApo150XOTIRFM, NA 1.45, Olympus) and an Electron-Multiplying Charge-Coupled Device camera (iXon^EM^+897-BI, Andor Technology). Fluorescence images of filaments labeled with Alexa Fluor 594 were merged with bright field images of cell bodies using ImageJ software version 1.54p (National Institutes of Health).

### Protein expression and purification

For expression of His-FlhA, *Salmonella* SJW1368 cells carrying pMKM130ES were grown in 2×YT medium supplemented with 100□μg□ml⁻¹ ampicillin at 30°C to an OD_600_ of 0.5–1.0, followed by incubation at 16°C for 24□h. Cells were harvested by centrifugation at 6,400□×□g for 10□min at 4°C. Cell pellets were resuspended in buffer containing 20□mM Tris-HCl (pH 8.0) and 3□mM EDTA and disrupted by sonication. Lysates were clarified by centrifugation at 20,000□×□g for 15□min at 4°C, and membrane fractions were collected by ultracentrifugation at 110,000□×□g for 1□h at 4°C.

Membranes were solubilized in 50□mM Tris-HCl (pH 8.0), 300□mM NaCl, 5% (w/v) glycerol, 20□mM imidazole, and 1% (w/v) LMNG at 4□°C for 1□h, followed by ultracentrifugation at 110,000□×□g for 1□h at 4°C to remove insoluble material. The supernatant was applied to a Ni-NTA agarose column (Qiagen) and washed with buffer containing 50□mM Tris-HCl (pH 8.0), 300□mM NaCl, 5% (w/v) glycerol, 20□mM imidazole, and 0.01% (w/v) LMNG. Bound proteins were eluted with a linear gradient of 100–400□mM imidazole.

Fractions containing His-FlhA were concentrated and further purified by size exclusion chromatography using a Superose 6 Increase 10/300 column (GE Healthcare) equilibrated in 50□mM Tris-HCl (pH 8.0), 150□mM NaCl, 1□mM EDTA, and 0.005% (w/v) LMNG. Peak fractions were pooled and concentrated to 2.5□mg□ml⁻¹, followed by buffer exchange into 50□mM piperazine-HCl (pH 5.5), 150□mM NaCl and 0.005% (w/v) LMNG.

For peptidisc reconstitution, peptidisc peptide (Peptidisc Biotech) was dissolved in 20□mM Tris-HCl (pH 7.8) at 5□mg□ml⁻¹ and mixed with purified His-FlhA at a 1:1 (w/w) ratio, followed by incubation for 30□min at room temperature. The sample was subjected to size exclusion chromatography using a Superose 6 Increase 10/300 column equilibrated in buffer containing 50□mM piperazine-HCl (pH 5.5) and 150□mM NaCl without detergent.

His-FlhA(Δ331-347) was expressed and purified as described for the wild-type protein, followed by peptidisc reconstitution at pH 5.5.

### pH-dependent reconstitution assay

To assess the effect of pH on FlhA ring formation, purified His-FlhA was concentrated to 2.5□mg□ml⁻¹ and buffer-exchanged using Amicon Ultra centrifugal filters. Buffer exchange was performed by dilution with an equal volume of buffer containing 50□mM buffering agent, 150□mM NaCl and 0.005% (w/v) LMNG, followed by concentration. This procedure was repeated four times.

The following buffers were used: 50 mM citric acid (pH 4.5 and 5.0), 50□mM piperazine-HCl (pH 5.5 and 11.0), 50□mM Bis-Tris-HCl (pH 6.0) and 50□mM Tris-HCl (pH 7.0 and 8.0), each containing 150□mM NaCl and 0.005% (w/v) LMNG.

Following buffer exchange, samples were subjected to peptidisc reconstitution as described previously. Briefly, peptidisc peptide was mixed with FlhA at a 1:1 (w/w) ratio, incubated for 30□min at room temperature, and analyzed by size exclusion chromatography in detergent-free buffer.

Ring formation was assessed by negative-stain electron microscopy before and after peptidisc reconstitution to evaluate pH-dependent assembly of FlhA under detergent-solubilized and membrane-mimetic conditions. Carbon-coated grids were glow-discharged using a JEC-3000FC sputter coater (JEOL, Japan). Purified FlhA samples (3 μl) were applied to the grids and blotted. The grids were then immediately stained with 3 μl of 2% (w/v) uranyl acetate solution and blotted. This staining step was repeated three times. After air-drying for 30 min, images were collected using a JEM-1400Flash transmission electron microscope (JEOL, Japan) operated at 80 kV.

### Cryo-EM sample preparation and data collection

Three different FlhA samples were vitrified under optimized grid preparation conditions for cryo-EM analysis. For wild-type FlhA, UltraAuFoil grids (R1.2/1.3, Au 200 mesh) were glow-discharged on the carbon side using a JEC-3000FC sputter coater at 20 mA for 20 s. A total of 3 μl of sample solution was applied to one side of the grid. Grids were prepared using a Vitrobot Mark IV (Thermo Fisher Scientific) equilibrated at 4 °C and 100% humidity. The grids were blotted for 3 s with a blot force of -10 and a drain time of 1 s, and then immediately plunged into liquid ethane.

For chemically crosslinked FlhA samples, epoxidized graphene grids^29^ were used. Immediately before vitrification, 0.3 μl of 0.5% (v/v) glutaraldehyde stock solution was added to 2.7 μl of sample solution to obtain a final glutaraldehyde concentration of 0.05% (v/v). After incubation for 15 s, the mixture was applied to one side of the grid. Grids were blotted for 1.5 s with a blot force of -25 and then immediately plunged into liquid ethane.

For the FlhA(Δ331–347) ring, Au-coated Quantifoil grids (R1.2/1.3, Cu 200 mesh) were glow-discharged on the carbon side using a JEC-3000FC sputter coater at 10 mA for 10 s. A total of 3 μl of sample solution was applied to one side of the grid. Grids were blotted for 3 s with a blot force of -10 and a drain time of 1 s before plunge-freezing in liquid ethane.

Excess ethane was removed by blotting with filter paper, and the grids were stored in liquid nitrogen. Cryo-EM image datasets were acquired using SerialEM ver. 4.0.29^30^ and yoneoLocr ver. 1.0.30^31^ on a JEM-Z300FSC (CRYO ARM™ 300, JEOL, Japan) operated at 300 kV, equipped with a K3 direct electron detector (Gatan, Inc.) in CDS mode. The Ω-type in-column energy filter was operated with a slit width of 20 eV for zero-loss imaging. The nominal magnification was x60,000, corresponding to a calibrated pixel size of 0.786 Å. Defocus values ranged from -0.5 to -2.0 μm. The total dose was 80 e^-^/Å² for the wild-type FlhA ring and 40 e^-^/Å² for the FlhA(Δ331–347) ring.

### Cryo-EM image processing of the wild-type FlhA ring

All image processing was performed using cryoSPARC v4.6.2^32^. The image processing procedure of the wild-type FlhA ring is described in Extended Data Fig. 7.

Cryo-EM data of the wild-type FlhA ring reconstituted in peptidisc were collected from three independent datasets comprising 1,276, 5,479 and 11,078 movies. A total of 17,833 movie stacks were subjected to motion correction and contrast transfer function (CTF) estimation, and micrographs with an estimated CTF resolution worse than 5□Å were discarded, leaving 15,581 micrographs for further analysis.

Particles were initially picked using Blob Picker (∼1.72 million particles). After removal of obvious false positives, particles were extracted (400 pixels, Fourier-cropped to 100 pixels) and subjected to multiple rounds of 2D classification, resulting in 514,707 particles.

These particles were used for *ab initio* reconstruction to generate four initial models, of which the best-resolved class (51.1%) was selected. Particles were then re-extracted (400 pixels, Fourier-cropped to 300 pixels) and refined by non-uniform refinement under C1 symmetry, yielding a reconstruction at 3.38□Å resolution from 292,355 particles.

To focus on the FlhA_C_ ring, density corresponding to the transmembrane region was removed by particle subtraction, followed by local refinement under C1 symmetry, volume re-centering and an additional round of local refinement.

For final refinement, particles were further classified and subjected to global and local CTF refinement, followed by non-uniform refinement under C9 symmetry, resulting in a reconstruction at 3.03□Å resolution from 274,285 particles.

### Cryo-EM image processing of the chemically crosslinked wild-type FlhA ring

To stabilize the FlhA ring structure reconstituted in peptidisc, we performed chemical crosslinking with glutaraldehyde. Then, image processing was performed as described above unless otherwise noted. A total of 11,643 movie stacks were processed, and 11,205 micrographs were retained after CTF-based filtering (<□5□Å) (Extended Data Fig. 10).

Particles were picked using Template Picker with 2D class averages of wild-type FlhA as templates (∼7.47 million particles), extracted (400 pixels, Fourier-cropped to 100 pixels), and subjected to multiple rounds of 2D classification, resulting in 828,437 particles.

Particles were re-extracted at a box size of 448 pixels and Fourier-cropped to 112 pixels, duplicate particles were removed, and further 2D classification yielded 781,196 particles for *ab initio* reconstruction. Four initial models were generated, and the best-resolved class (59.4%) was selected.

Selected particles were re-extracted (448 to 352 pixels) and refined by non-uniform refinement under C1 symmetry, yielding a reconstruction at 2.90□Å resolution from 464,081 particles.

Following particle subtraction to remove the transmembrane region, particles were refined under C1 symmetry with volume re-centering and local refinement. Final refinement included global and local CTF refinement and local refinement under C1 symmetry, yielding a reconstruction at 2.87□Å resolution, which was further refined under C9 symmetry to 2.60□Å resolution.

### Cryo-EM image processing of the FlhA(Δ331–347) ring

All data were processed using cryoSPARC v4.6.2^32^. The image processing procedure of the FlhA(Δ331–347) ring is described in Extended Data Fig. 8.

A total of 23,970 movie stacks were subjected to motion correction and CTF estimation, and micrographs with an estimated CTF resolution worse than 5□Å were discarded, leaving 22,738 micrographs for analysis.

Particles were picked using Template Picker (3,840,737 particles), extracted (448 pixels, Fourier-cropped to 112 pixels) and subjected to initial 2D classification using a randomly selected subset of 200,000 particles. A total of 123,249 particles from well-defined classes were selected for *ab initio* reconstruction followed by heterogeneous refinement.

The resulting subset (84,223 particles) was re-extracted (352 pixels) and refined under C1 symmetry, yielding a reconstruction at 3.73□Å resolution, which was subsequently used as a reference for heterogeneous refinement.

All particles were then classified by heterogeneous refinement using multiple reference volumes to improve separation of structural heterogeneity. Selected particles (1,136,535) were subjected to further 2D classification and non-uniform refinement, followed by global and local CTF refinement.

For high-resolution refinement, particles were re-extracted (352 pixels) and refined under C1 symmetry, yielding a reconstruction at 3.12□Å resolution from 845,751 particles. The same particles were subsequently refined under C9 symmetry to a final resolution of 2.73□Å.

To assess structural heterogeneity, particles were subjected to 3D classification followed by non-uniform refinement under C1 symmetry, yielding multiple classes with resolutions of 3.39–3.43□Å.

### Model building and refinement of the wild-type FlhA_C_ and FlhA(Δ331–347) rings

Initial models of the wild-type FlhA_C_ rings reconstructed with C1 and C9 symmetry were generated by fitting the crystal structure of FlhA_C_ (PDB ID: 3A5I) into the cryo-EM maps using UCSF ChimeraX^33^. The transmembrane region of FlhA(Δ331–347) was initially modeled using an AlphaFold3-predicted structure and subsequently rebuilt manually in Coot^34^. Models were iteratively adjusted by rigid-body fitting and local real-space refinement, with validation performed using MolProbity^35^. Final refinement against the cryo-EM maps was carried out in PHENIX^36^. Structural comparisons and figure preparation were performed using UCSF ChimeraX. Statistics for model refinement are summarized in Extended Data Tables 1 and 3.

## Notes

### Competing Interest Statement

The authors have declared no competing interest.

## References

1. Minamino, T. & Kinoshita, M. Structure, assembly, and function of flagella responsible for bacterial locomotion. EcoSal Plus eesp-0011-2023 (2023).

2. Minamino, T., Kinoshita, M. & Namba, K. Insight into distinct functional roles of the flagellar ATPase complex for flagellar assembly in *Salmonella*. Front. Microbiol. 13, 864178 (2022).

3. Minamino, T., Kinoshita, M., Morimoto, Y.V. & Namba, K. Activation mechanism of the bacterial flagellar dual-fuel protein export engine. Biophys. Physicobiol. 19, e190046 (2022).

4. Minamino, T., Morimoto, Y.V., Kinoshita, M. & Namba, K. Membrane voltage-dependent activation mechanism of the bacterial flagellar protein export apparatus. Proc. Natl. Acad. Sci. USA 118, e2026587118 (2021).

5. Minamino, T., Morimoto, Y.V., Hara, N., Aldridge, P.D. & Namba, K. The bacterial flagellar type III export gate complex is a dual fuel engine that can use both H^+^ and Na^+^ for flagellar protein export. PLoS Pathog. 12, e1005495 (2016).

6. Minamino, T., Kinoshita, M., Morimoto, Y.V. & Namba K. The FlgN chaperone activates the Na^+^-driven engine of the *Salmonella* flagellar protein export apparatus. *Commun*. Biol. 4, 335 (2021).

7. Bange, G., et al. FlhA provides the adaptor for coordinated delivery of late flagella building blocks to the type III secretion system. Proc. Natl. Acad. Sci. USA 107, 11295–11300 (2010).

8. Minamino, T., et al. Interaction of a bacterial flagellar chaperone FlgN with FlhA is required for efficient export of its cognate substrates. Mol. Microbiol. 83, 775–788 (2012).

9. Abrusci, P., et al. Architecture of the major component of the type III secretion system export apparatus. Nat. Struct. Mol. Biol. 20, 99–104 (2013).

10. Kinoshita, M., Hara, N., Imada, K., Namba, K. & Minamino, T. Interactions of bacterial chaperone-substrate complexes with FlhA contribute to co-ordinating assembly of the flagellar filament. Mol. Microbiol. 90, 1249–1261 (2013).

11. Xing, Q., et al. Structure of chaperone-substrate complexes docked onto the export gate in a type III secretion system. Nat. Commun. 9, 1773 (2018).

12. Minamino, T., Morimoto, Y.V., Hara, N. & Namba, K. An energy transduction mechanism used in bacterial type III protein export. Nat. Commun. 2, 475 (2011).

13. Terahara, N. et al. Insight into structural remodeling of the FlhA ring responsible for bacterial flagellar type III protein export. Sci. Adv. 4, eaao7054 (2018).

14. Inoue, Y., et al. The FlhA linker mediates flagellar protein export switching during flagellar assembly. Commun. Biol. 4, 646 (2021).

15. Erhardt, M., et al. Mechanism of type-III protein secretion: Regulation of FlhA conformation by a functionally critical charged-residue cluster. Mol. Microbiol. 104, 234– 249 (2017).

16. Kuhlen, L., et al. Structure of the core of the type III secretion system export apparatus. Nat. Struct. Mol. Biol. 25, 583–590 (2018).

17. Kuhlen, L., et al. The substrate specificity switch FlhB assembles onto the export gate to regulate type three secretion. Nat. Commun. 11, 1296 (2020).

18. Kuhlen, L., Johnson, S., Cao, J., Deme, J.C. & Lea, S.M. Nonameric structures of the cytoplasmic domain of FlhA and SctV in the context of the full-length protein. PLoS One 16, e0252800 (2021).

19. Kinoshita, M., et al. A β-cap on the FliPQR protein-export channel acts as the cap for initial flagellar rod assembly. Proc. Natl. Acad. Sci. USA 122, e2507221122 (2025).

20. Kawamoto, A., et al. Common and distinct structural features of *Salmonella* injectisome and flagellar basal body. Sci. Rep. 3, 3369 (2013).

21. Minamino, T. & Namba, K. Distinct roles of the FliI ATPase and proton motive force in bacterial flagellar protein export. Nature 451, 485–488 (2008).

22. Kinoshita, M., Minamino, T., Uchihashi, T. & Namba, K. FliH and FliI help FlhA bring strict order to flagellar protein export in *Salmonella*. Commun. Biol. 7, 366 (2024).

23. Einenkel, R. et al. The FliI ATPase couples ATP hydrolysis to substrate switching in bacterial flagellar type-III secretion. mBio 17, e0235425 (2026).

24. Saijo-Hamano, Y., Minamino, T., Macnab, R.M. & Namba, K. Structural and functional analysis of the C-terminal cytoplasmic domain of FlhA, an integral membrane component of the type III flagellar protein export apparatus in *Salmonella*. J. Mol. Biol. 343, 457–466 (2004).

25. Carlson, M.L., et al. The Peptidisc, a simple method for stabilizing membrane proteins in detergent-free solution. eLife 7, e34085 (2018).

26. McMurry, J.L., Van Arnam, J.S., Kihara, M. & Macnab, R.M. Analysis of the cytoplasmic domains of Salmonella FlhA and interactions with components of the flagellar export machinery. J. Bacteriol. 186, 7586–7592 (2003).

27. Hara, N., Namba, K. & Minamino, T. Genetic characterization of conserved charged residues in the bacterial flagellar type III export protein FlhA. PLoS ONE 6, e22417 (2011).

28. Barker, C.S, Inoue, T., Meshcheryakova, I.V., Kitanobo, S. & Samatey, F.A. Function of the conserved FHIPEP domain of the flagellar type III export apparatus, protein FlhA. Mol. Microbiol. 100, 278–288 (2016).

## References

29. Fujita, J. et al. Epoxidized graphene grid for highly efficient high-resolution cryoEM structural analysis. Sci. Rep. 13, 2279 (2023).

30. Mastronarde, D.N. Automated electron microscope tomography using robust prediction of specimen movements. J. Struct. Biol. 152, 36–51 (2005).

31. Yonekura, K., Maki-Yonekura, S., Naitow, H., Hamaguchi, T. & Takaba, K. Machine learning-based real-time object locator/evaluator for cryo-EM data collection. Commun. Biol. 4, 1044 (2021).

32. Punjani, A., Rubinstein, J.L., Fleet, D.J. & Brubaker, M.A. cryoSPARC: algorithms for rapid unsupervised cryo-EM structure determination. Nat. Methods 14, 290–296 (2017).

33. Pettersen, E., et al. UCSF ChimeraX: Structure visualization for researchers, educators, and developers. Protein Sci. 30, 70–82 (2021).

34. Emsley, P., Lohkamp, B., Scott, W.G. & Cowtan, K. Features and development of *Coot*. Acta Crystallogr. D Struct. Biol. 66, 486–501 (2010).

35. Chen, V.B. MolProbity: all-atom structure validation for macromolecular crystallography. Acta Crystallogr. D Biol. Crystallogr. 66, 12–21 (2010).

36. Liebschner, D., et al. Macromolecular structure determination using X-rays, neutrons and electrons: recent developments in Phenix. Acta Crystallogr. D Struct. Biol. 75, 861–877 (2019).

