## Supplementary Information for "Structure of an active FlhA ring reveals allosteric coupling between substrate specificity and export engine activation"

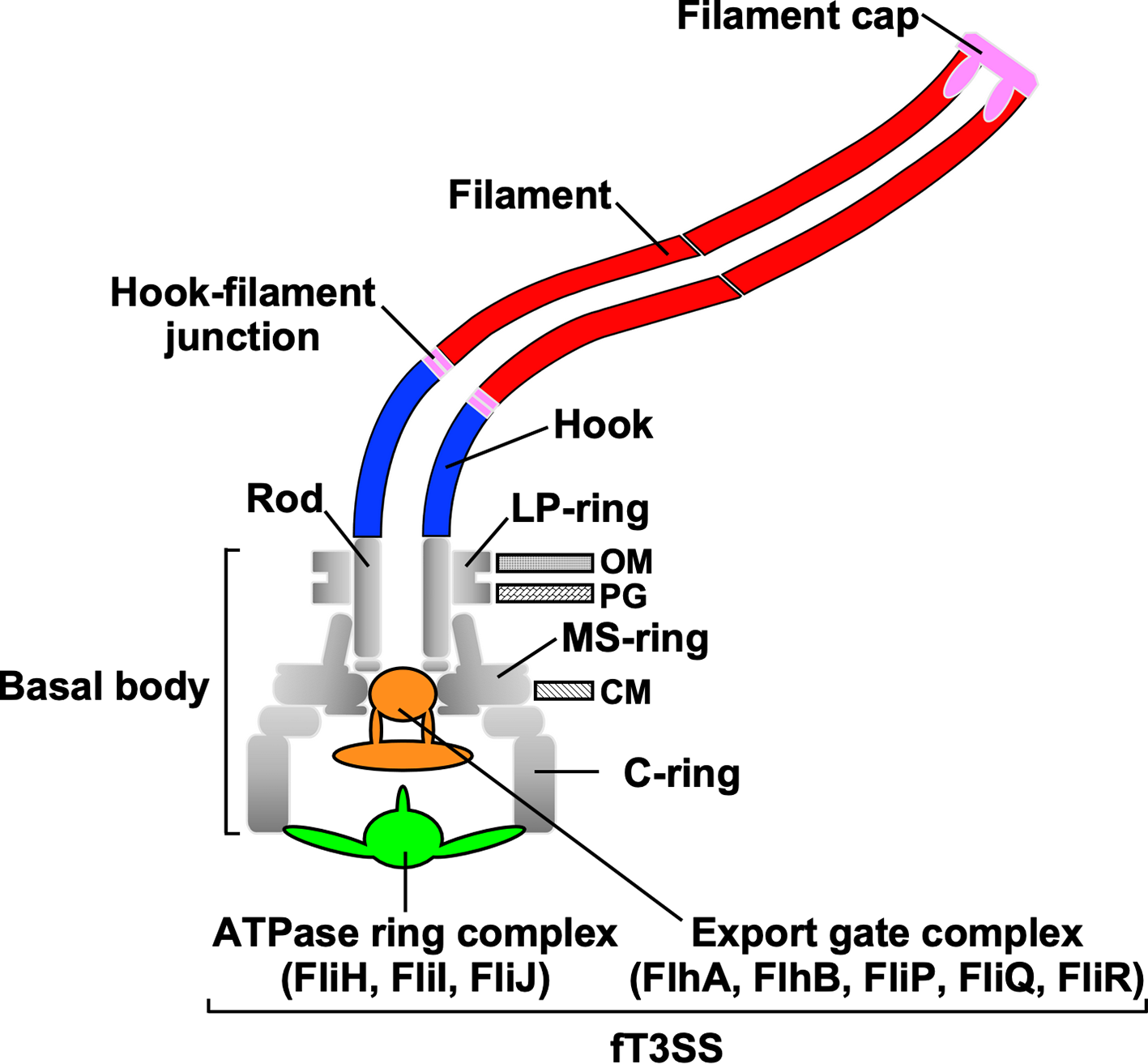

**Extended Data Fig. 1. Schematic diagram of the bacterial flagellum.** The bacterial flagellum comprises a basal body with MS, C, and LP rings, and an axial structure consisting of the rod, hook, hook-filament junction, filament, and filament cap. Assembly of the axial structure beyond the cellular membranes is driven by the flagellar type III secretion system (fT3SS), located at the base of the flagellum, which exports axial component proteins from the cytoplasm to the distal end of the growing structure. The fT3SS consists of a transmembrane export gate complex (FlhA, FlhB, FliP, FliQ, and FliR) and a cytoplasmic ATPase ring complex (FliH, FliI, and FliJ).

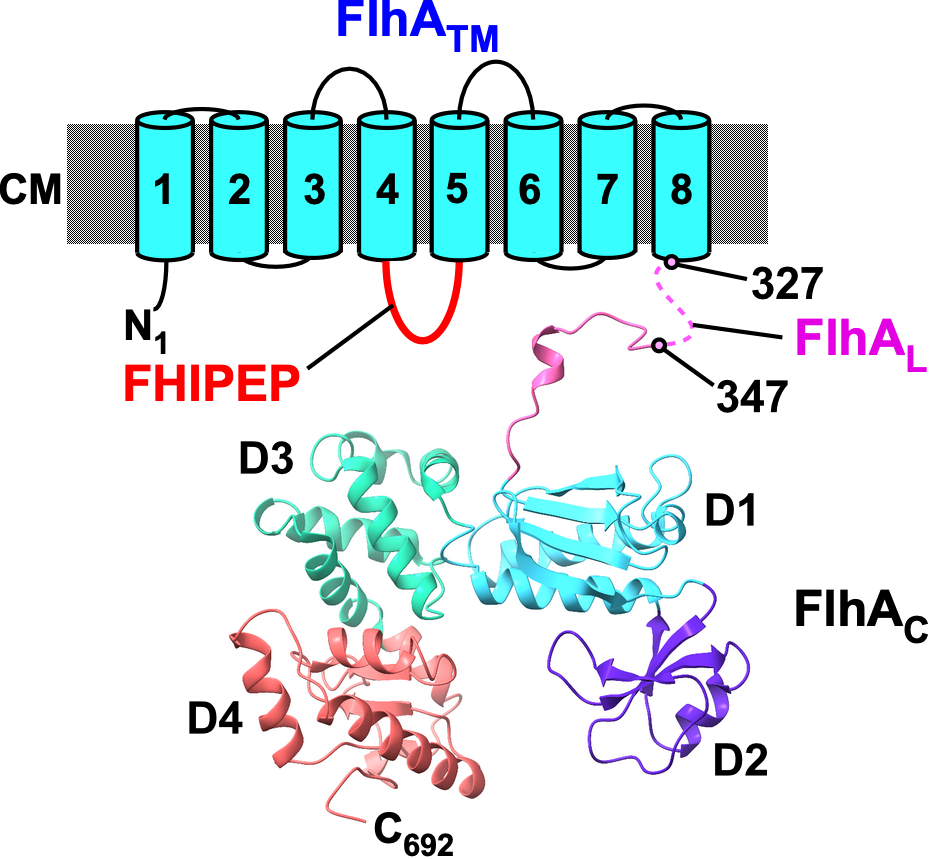

**Extended Data Fig. 2. Domain organization of FlhA.** *Salmonella* FlhA (UniProt ID: P40729) is a 692-amino-acid protein comprising three regions: an N-terminal transmembrane domain (FlhA_TM_, residues 1–327) with eight predicted transmembrane helices (TM1–TM8), a C-terminal cytoplasmic domain (FlhA_C_, residues 362–692), and a flexible linker (FlhA_L_, residues 328–361) connecting FlhA_TM_ and FlhA_C_. The cytoplasmic domain (residues 327–692) has been solved by X-ray crystallography (PDB ID: 3A5I) and consists of four subdomains (D1–D4). Residues 328–346 within FlhA_L_ (dashed line) are not resolved in this structure owing to conformational flexibility. A highly conserved cytoplasmic loop between transmembrane helices TM4 and TM5, termed the FHIPEP (flagellum/hypersensitive response/invasion protein export protein) domain (residues 141–208), is essential for H⁺-coupled protein export and substrate specificity switching.

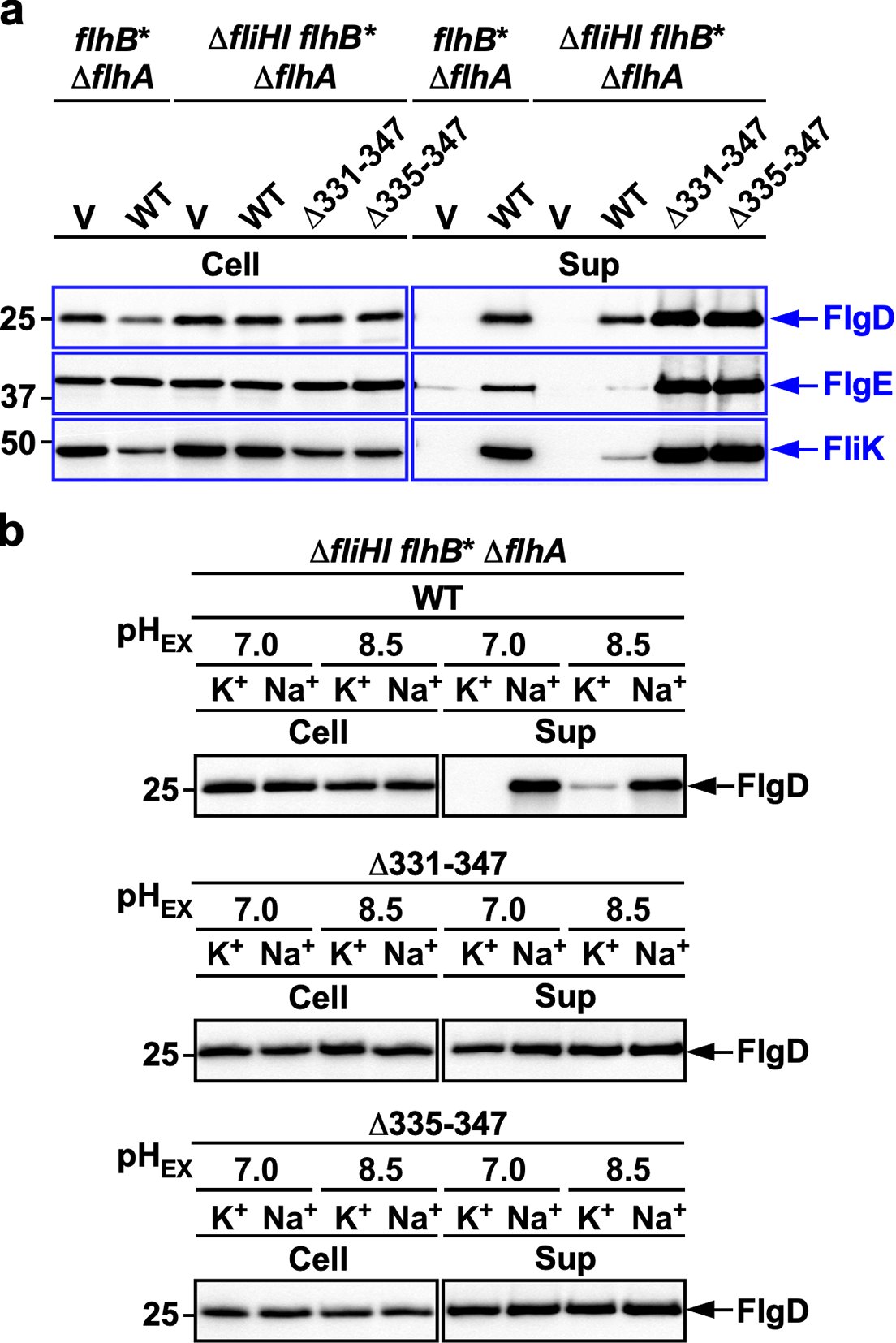

**Extended Data Fig. 3**. **Effect of *flhA* linker deletions on protein secretion efficiency in the Δ*fliH-fliI flhB*(P28T) mutant. (a)** Whole-cell (Cell) and culture supernatant (Sup) fractions were prepared from *Salmonella* NH001 (Δ*flhA*) carrying pTrc99AFF4 (V) or pMM130 (WT), and NH004 (Δ*fliHI flhB** Δ*flhA*) carrying pTrc99AFF4 (V), pMM130 (WT), pMKM130-207 (Δ331–347) or pMKM130-212 (Δ335–347), and analyzed by immunoblotting with the indicated polyclonal antibodies. Molecular mass markers (kDa) are indicated. Data are representative of three independent experiments. **(b)** Na^+^- and membrane potential-dependent FlgD secretion in the absence of FliH and FliI. Immunoblots of whole-cell (Cell) and supernatant fractions from *Salmonella* NH004 carrying pMM130 (top), pMKM130-207 (middle) or pMKM130-212 (bottom), grown at 30 °C in T-broth at pH 7.0 or 8.5 with either 100 mM K⁺ or Na⁺. Molecular mass markers (kDa) are indicated. Data are representative of three independent experiments.

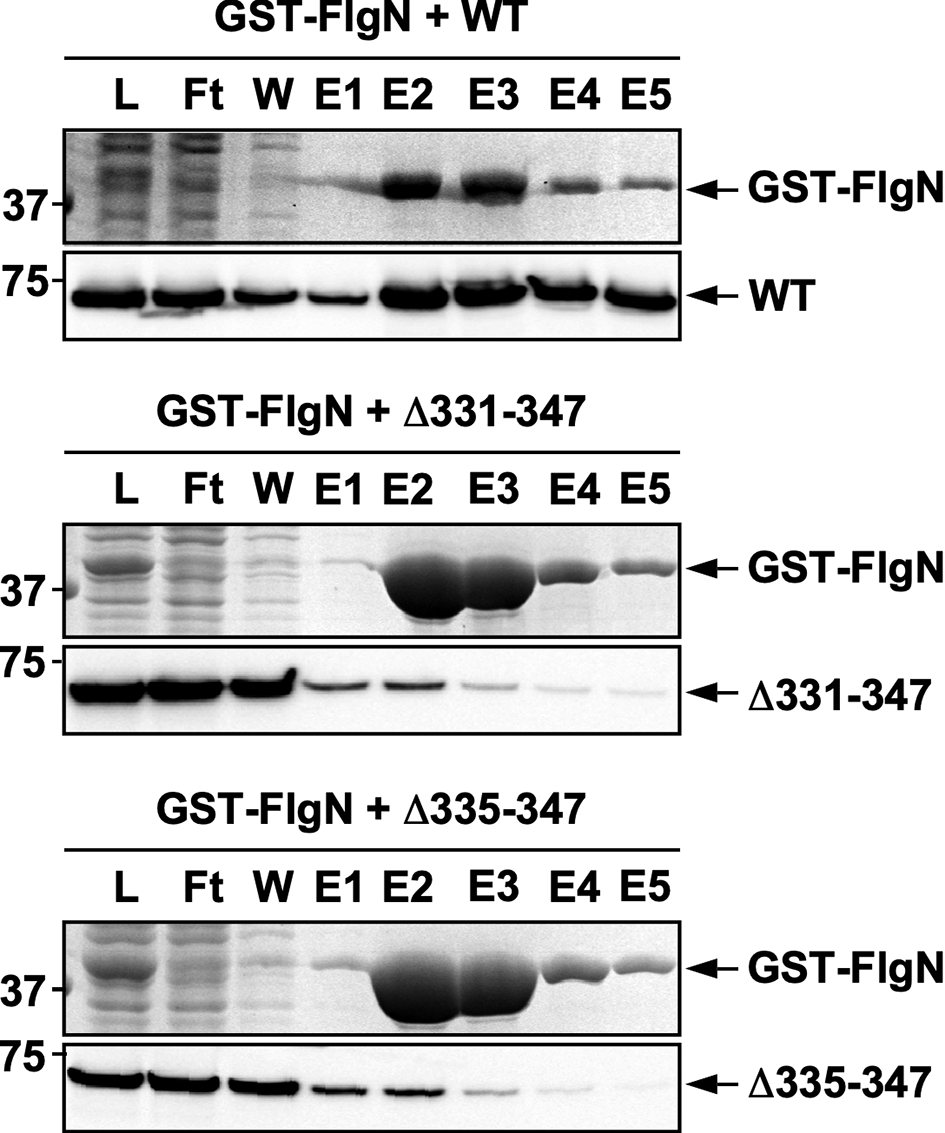

**Extended Data Fig. 4. Effect of FlhA linker deletions on interaction between FlhA and the filament-type export chaperone FlgN.** Soluble fractions (L) prepared from the Salmonella SJW1368 (Δ*cheW-flhD*) strain expressing GST-FlgN were mixed with cell lysates from the *Salmonella* NH003 (Δ*fliH-fliI* *flhA*) strain carrying pMM130 (WT, upper panels), pMKM130-207 (Δ331-347, middle panels), or pMKM130-212 (Δ335-347, lower panels). The mixtures were loaded onto a GST column. After washing with PBS, bound proteins were eluted with buffer containing 10 mM reduced glutathione. The eluted fractions were analyzed by CBB staining for GST-FlgN and immunoblotting with a polyclonal anti-FlhA_C_ antibody.

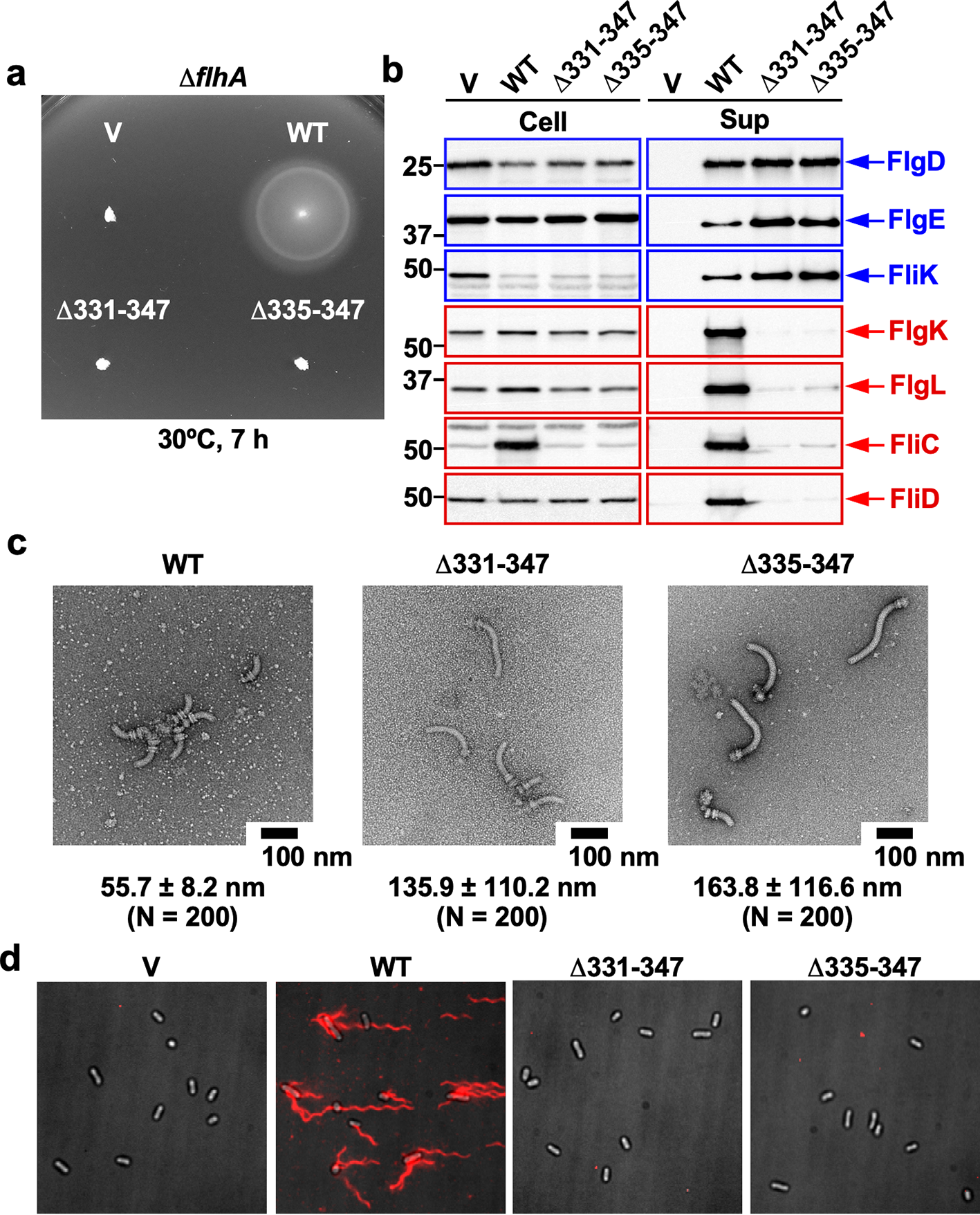

**Extended Data Fig. 5. Characterization of *flhA* linker deletion mutants in the presence of FliH and FliI. (a)** Soft-agar motility of *Salmonella* NH001 (Δ*flhA*) carrying pTrc99AFF4 (V), pMM130 (WT), pMKM130-207 (Δ331–347) or pMKM130-212 (Δ335–347). Plates were incubated at 30°C for 7 h. Data are representative of at least seven independent experiments. **(b)** Secretion of FlgD, FlgE, FliK, FlgK, FlgL FliC and FliD. Whole-cell (Cell) and supernatant (Sup) fractions were prepared from the transformants shown in (a) and analyzed by immunoblotting with the indicated polyclonal antibodies. Hook- and filament-type substrates are shown in blue and red, respectively. Molecular mass markers (kDa) are indicated. Data are representative of at least three independent experiments. **(c)** Electron micrographs of hook-basal bodies isolated from the transformants shown in (a). Mean hook length ± s.d. are shown; *N* indicates the number of structures analyzed. **(d)** Fluorescence imaging of the transformants shown in (a). Cells were grown from fresh colonies in L-broth containing ampicillin to stationary phase, and flagellar filaments were labelled with Alexa Fluor 594. Fluorescence images (red) were merged with corresponding bright-field images.

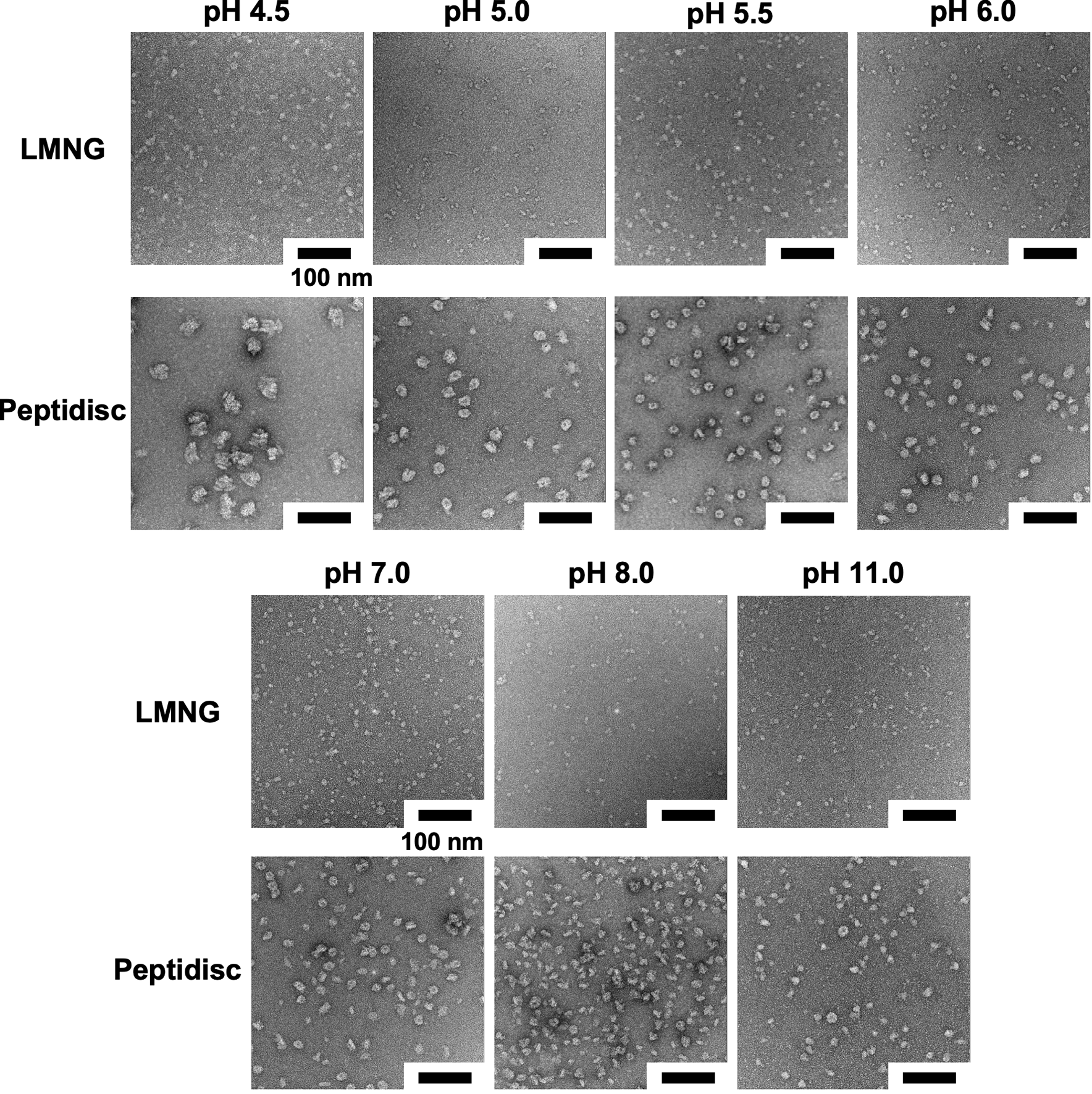

**Extended Data Fig. 6. Effect of pH on FlhA ring formation.** Purified wild-type FlhA solubilized in LMNG was concentrated to 2.5 mg ml⁻¹ and buffer-exchanged using Amicon Ultra centrifugal filters. The following buffers were used: 50 mM citric acid (pH 4.5 and 5.0), 50 mM piperazine-HCl (pH 5.5 and 11.0), 50 mM Bis-Tris-HCl (pH 6.0), and 50 mM Tris-HCl (pH 7.0 and 8.0), each containing 150 mM NaCl and 0.005% (w/v) LMNG. LMNG was then replaced by peptidisc under each pH condition. Scale bar, 100 nm.

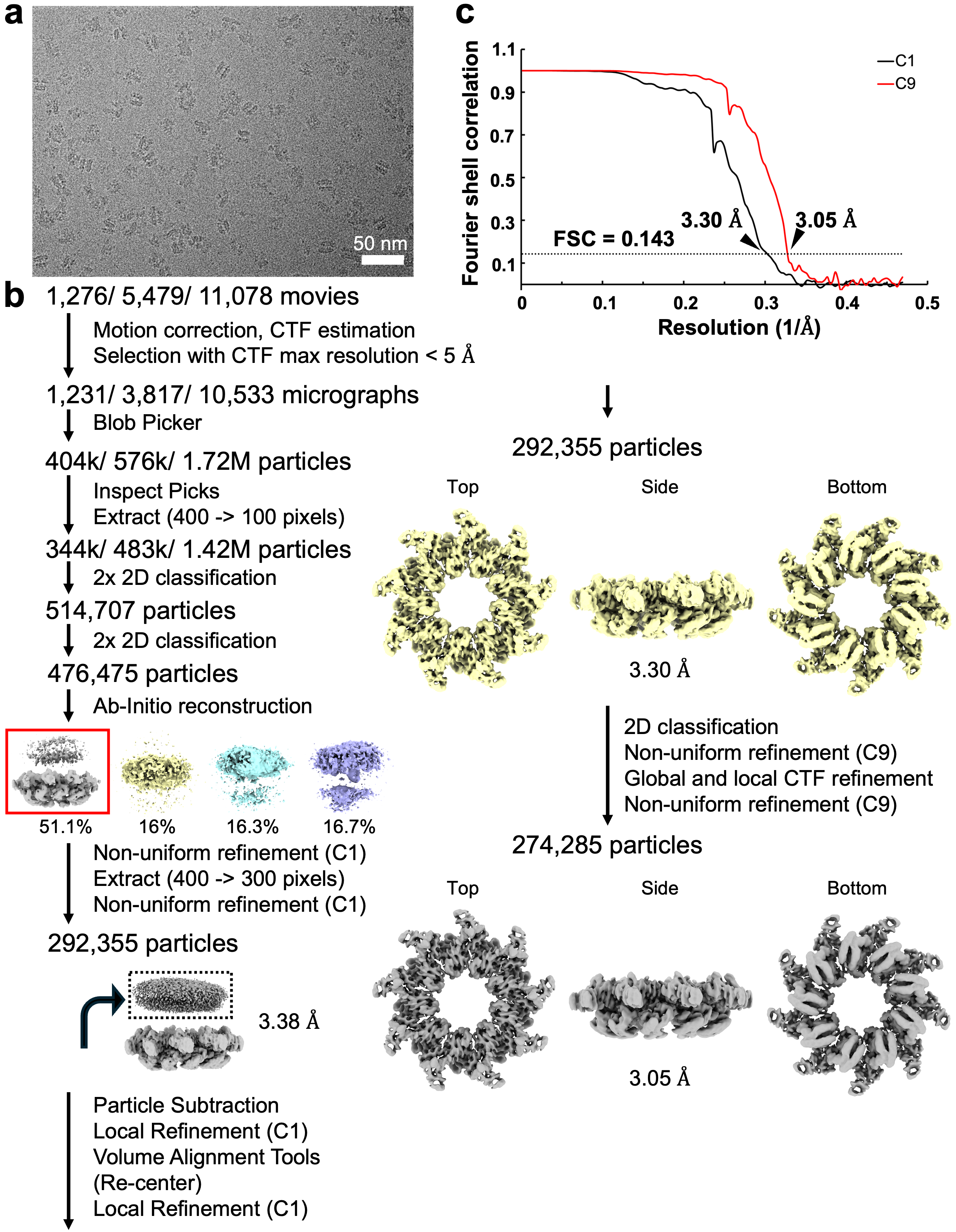
**Extended Data Fig. 7**. **Cryo-EM data processing workflow of the wild-type FlhA_C_ ring. (a)** Representative cryo-EM micrograph of the wild-type FlhA ring complex reconstituted in peptidisc. **(b)** CryoSPARC processing workflow. **(c)** Fourier shell correlation (FSC) curves of the final density maps of the FlhA_C_ ring refined with either C1 (black line, EMD-80484) or C9 symmetry (red line, EMD-80485).

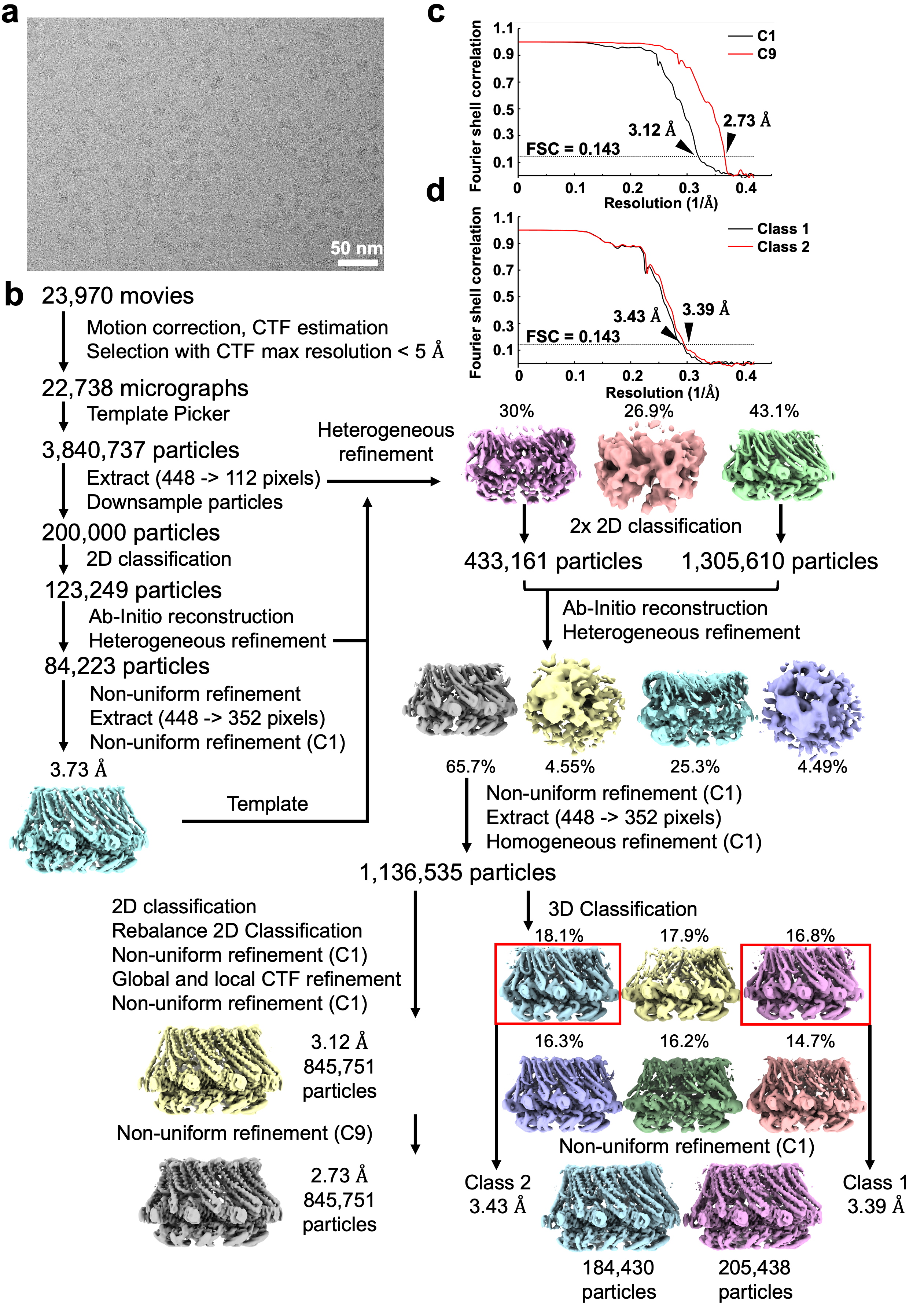
**Extended Data Fig. 8**. **Cryo-EM data processing workflow of the FlhA(Δ331–347　 ring. (a)** Representative cryo-EM micrograph of the FlhA(Δ331–347) ring reconstituted in peptidisc. **(b)** CryoSPARC processing workflow. **(c)** Fourier shell correlation (FSC) curves of the final density maps of the FlhA(Δ331–347) ring refined with either C1 (black line) or C9 symmetry (red line, EMD-80488). **(d)** Fourier shell correlation (FSC) curves of the final density maps of the two classes of the FlhA(Δ331–347) ring refined with C1 (class 1, EMD-80489; class 2, EMD-80490).

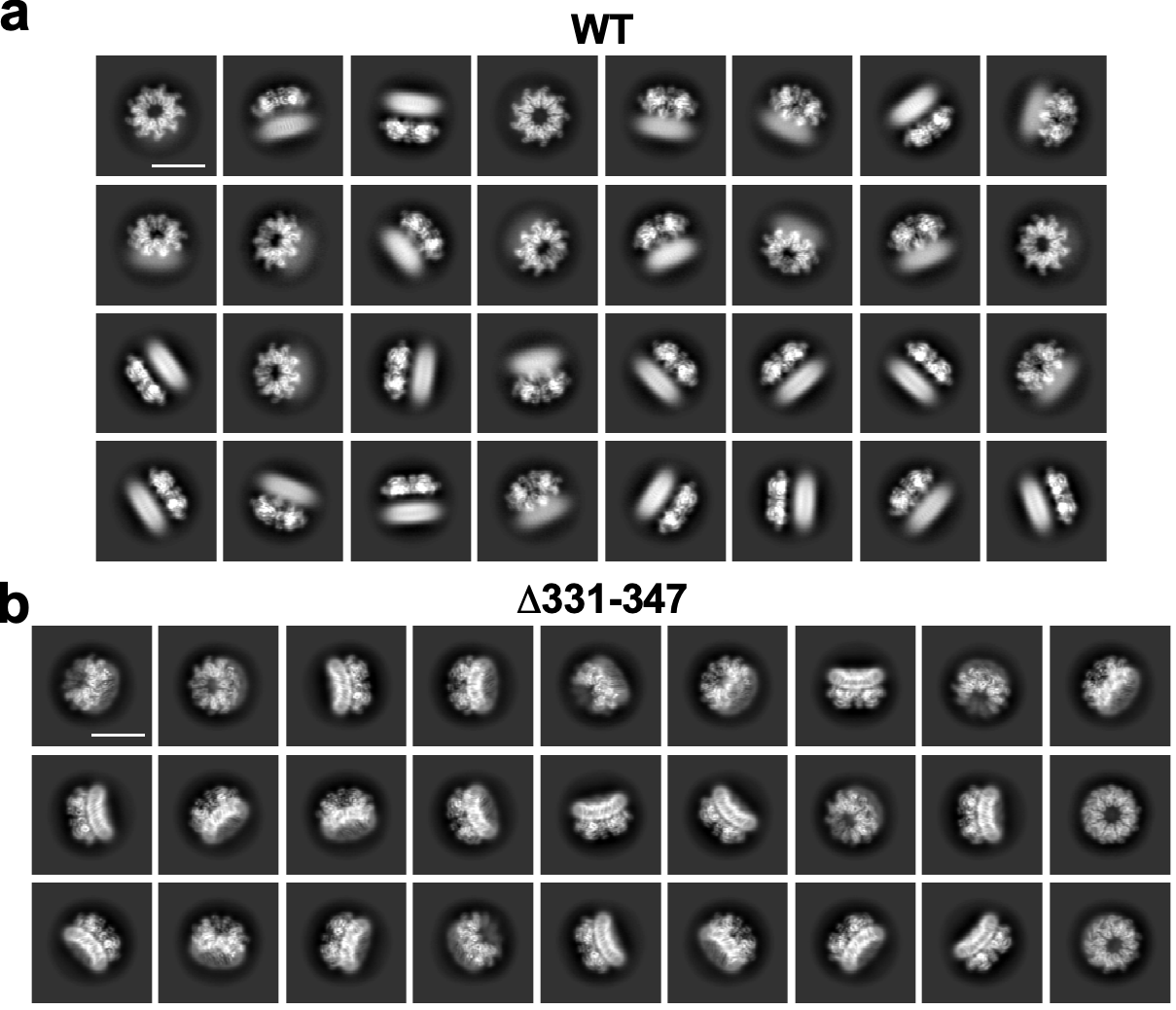

**Extended Data Fig. 9**. **Representative 2D class averages of FlhA ring complex. (a)** Wild-type FlhA ring**. (b)** FlhA(Δ331–347) ring. 2D class averages were obtained by single particle cryo-EM image analysis. Scale bar, 15 nm.

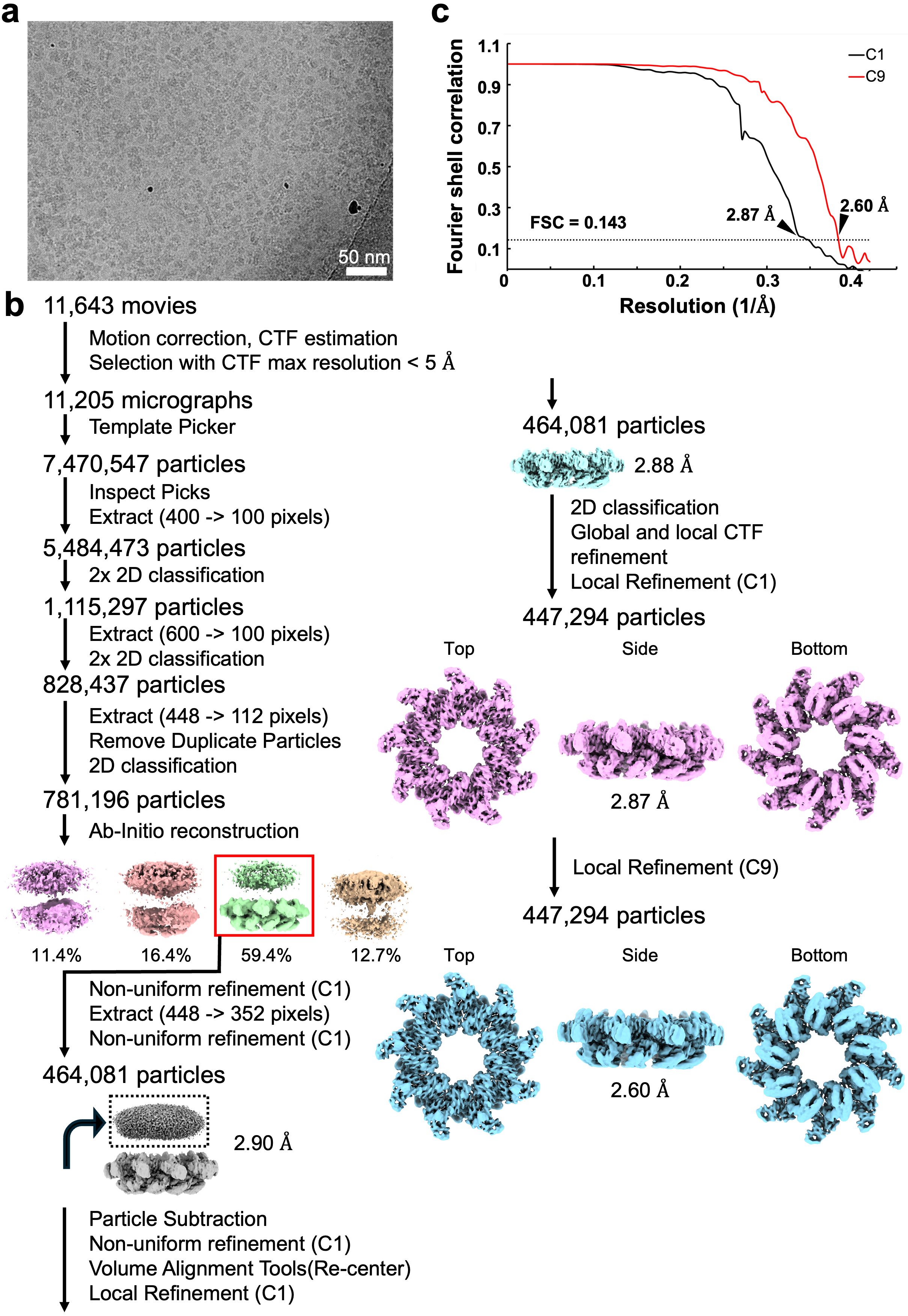
**Extended Data Fig. 10**. **Cryo-EM data processing workflow of the chemical-crosslinked wild-type FlhA_C_ ring. (a)** Representative cryo-EM micrograph of the cross-linked FlhA ring reconstituted in peptidisc. **(b)** CryoSPARC processing workflow. **(c)** Fourier shell correlation (FSC) curves of the final density maps of the cross-linked FlhA_C_ ring refined with either C1 (black line, EMD-80486) or C9 symmetry (red line, EMD-80487).

**Extended Data Fig. 11. Cryo-EM map of the wild-type FlhA ring reconstructed with C9 symmetry. (a)** Cryo-EM density map of the wild-type FlhA ring. Density corresponding to FlhA_L_ becomes visible at a higher density threshold. The linker adopts an extended conformation that spatially separates the FlhA_C_ ring from the FlhA_TM_ ring. **(b)** Fourier shell correlation (FSC) curve of the final density map of the wild-type FlhA ring reconstructed with C9 symmetry.

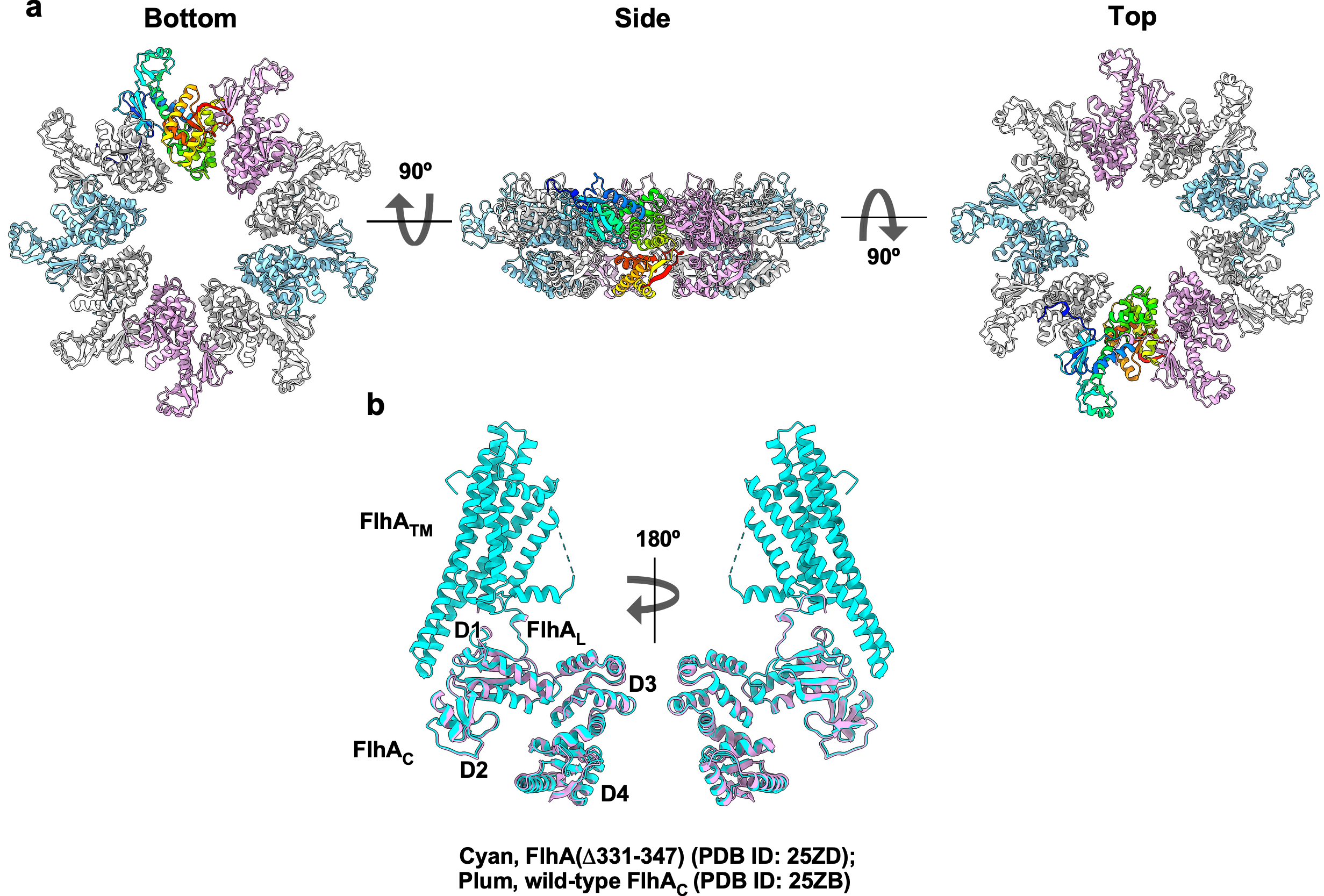
**Extended Data Fig. 12**. **Cryo-EM structure of the chemically crosslinked FlhA_C_ ring (a)** Cα ribbon diagram of the atomic model of the FlhA_C_ ring complex (PDB ID: 25ZB). **(b)** Structural comparison between wild-type FlhA_C_ and FlhA(Δ331–347), showing high structural similarity. Domain D1 of wild-type FlhA_C_ was superimposed onto that of FlhA(Δ331–347) (PDB ID: 25ZB), yielding an RMSD of 1.146 Å, calculated using ChimeraX.

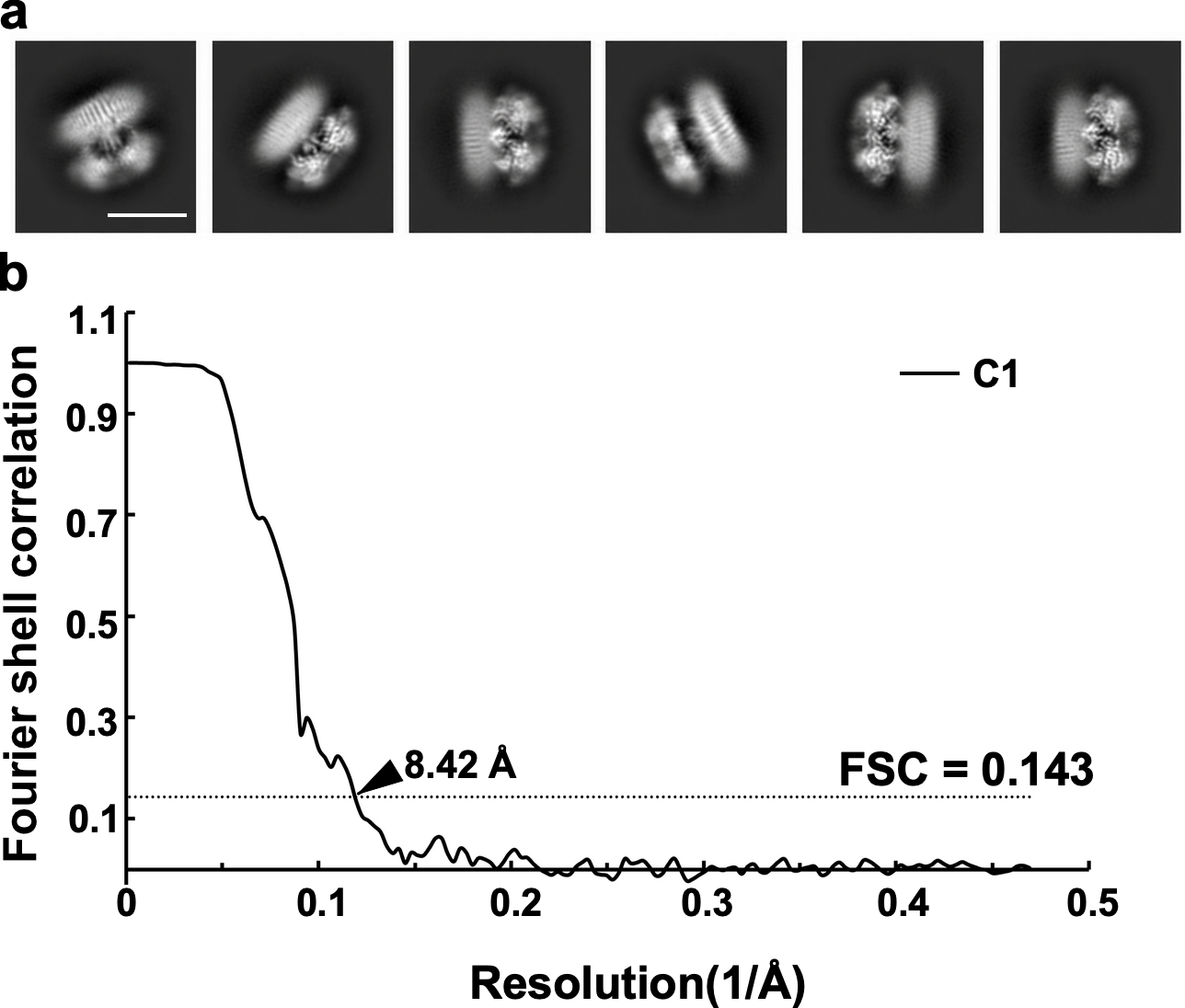

**Extended Data Fig. 13**. **Cryo-EM image analysis of the wild-type FlhA ring (a)** Selected 2D class averages of the wild-type FlhA ring exhibiting prominent FHIPEP-like structural features extending from the center of the FlhA_TM_ ring. Scale bar, 13 nm. **(b)** Fourier shell correlation (FSC) curve of the final density map of the crosslinked FlhA ring refined with C1 symmetry.

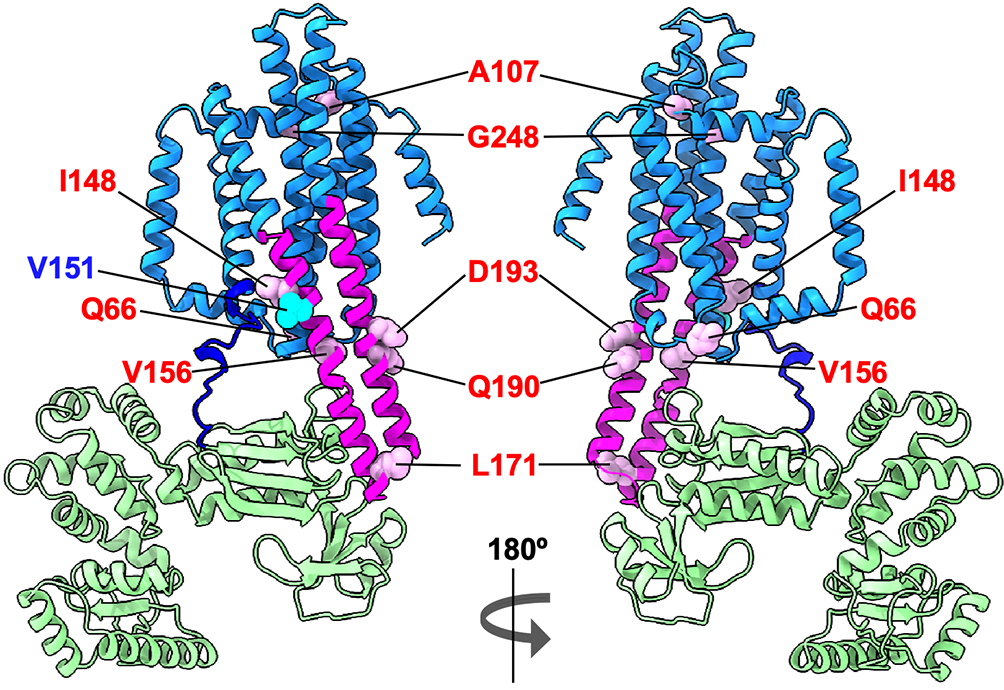

**Extended Data Fig. 14**. **Mapping of the *flhA*(V151) mutation site and intragenic suppressor mutations onto the FlhA(Δ331–347) structure (PDB ID: 25ZG).** FlhA_TM_, dogger blue; FlhA_L_, blue; FlhA_C_, light green. The *flhA*(V151L) mutation site is shown in cyan, and intragenic suppressor mutation sites are shown in plum. Val151 forms an intramolecular hydrophobic contact with Val350 in FlhA_L_ (see Fig. 3). The *flhA*(V151L) mutation disrupts the ordered secretion of hook-type export substrates and inhibits substrate specificity switching, resulting in a non-motile phenotype on soft agar plates. Intragenic suppressor mutations, including Q66R, A107P, A107T, A107V, I148T, V156A, V156M, L171I, Q190L, D193E and G248S, restore motility of the *flhA*(V151L) mutant. Residues I148, V151, V156, L171, Q190 and D193 are located within the FHIPEP domain. Q66 is located in a cytoplasmic loop connecting TM2 and TM3, whereas A107 is located in a periplasmic loop connecting TM3 and TM4. G248 is located within TM6. These genetic data suggest that the V151L substitution induces conformational rearrangements in both FlhA_TM_ and the FHIPEP domain, thereby locking the fT3SS in the hook-type secretion state. Intragenic suppressor mutations restore the ability of the fT3SS to switch substrate specificity from hook-type to filament-type export. Together, these data support a model in which topological remodeling of FlhA_TM_ is coupled to substrate specificity switching in the fT3SS.

Extended Data Table 1. CryoEM data collection, processing, and refinement statistics of the FlhA(Δ331–347) ring structure.

| Dataset | FlhA(Δ331–347) in C9 symmetry | FlhA(Δ331–347) in C1 symmetry  Class 1 | FlhA(Δ331–347) in C1 symmetry  Class 2 |
| --- | --- | --- | --- |
| EMDB  PDB | EMD-80488  25ZD | EMD-80489  25ZG | EMD-80490  25ZI |
| **Data collection and processing** | | | |
| Magnification | 60,000 | 60,000 | 60,000 |
| Voltage (kV) | 300 | 300 | 300 |
| Electron exposure (e−/Å^2^) | 40 | 40 | 40 |
| Defocus range (μm) | -1.8 – -2.2 | -1.8 – -2.2 | -1.8 – -2.2 |
| Pixel size (Å) | 0.786 | 0.786 | 0.786 |
| Symmetry imposed | C9 | C1 | C1 |
| Imported movies (no.) | 23,970 | 23,970 | 23,970 |
| Initial particle images (no.) | 3,840,737 | 3,840,737 | 3,840,737 |
| Final particle images (no.) | 845,751 | 205,438 | 184,430 |
| Map resolution (Å) | 2.73 | 3.39 | 3.43 |
| FSC threshold | 0.143 | 0.143 | 0.143 |
| **Refinement** | | | |
| Initial model used (PDB code) |  | 3A5I | 3A5I |
| Model resolution (Å) | 2.7/3.0/4.5 | 2.8/2.9/3.1 | 3.4/3.4/3.8 |
| FSC threshold | 0/0.143/0.5 | 0/0.143/0.5 | 0/0.143/0.5 |
| Model vs. Data CC (mask) | 0.68 | 0.82 | 0.82 |
| (volume) | 0.69 | 0.82 | 0.81 |
| **Model composition** | | | |
| Non-hydrogen atoms | 42,606 | 43,939 | 43,745 |
| Protein residues | 5,589 | 5,758 | 5,785 |
| Ligands | 0 | 0 | 0 |
| **Validation** | | | |
| Bond length (Å) | 0.001 | 0.002 | 0.003 |
| Bond angles (^ο^) | 0.348 | 0.421 | 0.447 |
| MolProbity score | 1.21 | 1.48 | 1.54 |
| Clash score | 4.35 | 6.29 | 7.00 |
| Rotamer outliers (%) | 0.40 | 1.43 | 1.51 |
| **Ramachandran plot** | | | |
| Favored (%) | 99.03 | 97.95 | 97.92 |
| Allowed (%) | 0.97 | 2.05 | 2.08 |
| Disallowed (%) | 0 | 0 | 0 |

**Extended Data Table 2. Locations of the secondary structures in FlhA transmembrane domains in the assembled FlhA(Δ331–347) ring complex.**

| **FlhA(Δ331–347) in C9 symmetry (PDB ID: 25ZD)** | | | | | | | | | | |
| --- | --- | --- | --- | --- | --- | --- | --- | --- | --- | --- |
|  |  | **Chains (residues)** | | | | | | | | |
|  |  | **A–I** | | | | | | | | |
| **α1** | **TM1** | – | | | | | | | | |
| **α2** | **TM2** | 41–63 | | | | | | | | |
| **α3** | **TM3** | 68–102 | | | | | | | | |
| **α4** |  | 109–119 | | | | | | | | |
| **α5** | **TM4 + FHIPEP** | 123–173 | | | | | | | | |
| **α6** | **TM5 + FHIPEP** | 178–230 | | | | | | | | |
| **α7** | **TM6** | 235–273 | | | | | | | | |
| **α8** |  | 277–287 | | | | | | | | |
| **α9** | **TM7** | – | | | | | | | | |
| **α10** | **TM8** | 312–330 | | | | | | | | |
| **FlhA(Δ331–347) in C1 symmetry (PDB ID: 25ZG)**  **Class 1** | | | | | | | | | | |
|  |  | **Chains (residues)** | | | | | | | | |
|  |  | **A** | **B** | **C** | **D** | **E** | **F** | **G** | **H** | **I** |
| **α1** | **TM1** | 25–34 | – | – | 25–36 | 27–35 | – | 30–35 | 28–31 | – |
| **α2** | **TM2** | 41–63 | 41–63 | 41–63 | 41–63 | 41–63 | 41–63 | 41–63 | 41–63 | 41–63 |
| **α3** | **TM3** | 69–102 | 69–99 | 68–99 | 68–102 | 68–102 | 68–102 | 69–102 | 68–102 | 68–102 |
| **α4** |  | 109–119 | 109–119 | 109–119 | 109–119 | 108–119 | 109–119 | 109–119 | 109–119 | 109–119 |
| **α5** | **TM4 + FHIPEP** | 123–173 | 123–173 | 123–173 | 123–173 | 123–173 | 123–173 | 123–173 | 123–173 | 123–173 |
| **α6** | **TM5 + FHIPEP** | 178–229 | 178–229 | 178–229 | 178–230 | 178–230 | 178–229 | 178–229 | 178–229 | 178–229 |
| **α7** | **TM6** | 235–273 | 235–272 | 235–273 | 235–272 | 235–272 | 235–272 | 235–272 | 235–273 | 235–273 |
| **α8** |  | 277–287 | 277–287 | 277–287 | 277–287 | 277–287 | 277–287 | 277–287 | 277–287 | 277–287 |
| **α9** | **TM7** | 289–304 | 289–304 | 300–304 | 289–304 | 289–304 | 289–304 | 289–304 | 289–304 | 289–304 |
| **α10** | **TM8** | 312–330 | 311–330 | 311–330 | 312–330 | 312–330 | 312–330 | 312–330 | 312–330 | 312–330 |
| **FlhA(Δ331–347) in C1 symmetry (PDB ID: 25ZI)**  **Class 2** | | | | | | | | | | |
|  |  | **Chains (residues)** | | | | | | | | |
|  |  | **A** | **B** | **C** | **D** | **E** | **F** | **G** | **H** | **I** |
| **α1** | **TM1** | 25–34 | – | – | – | 27–35 | – | – | – | – |
| **α2** | **TM2** | 41–63 | 41–63 | 41–63 | 41–63 | 41–63 | 41–63 | 41–63 | 41–63 | 41–63 |
| **α3** | **TM3** | 69–102 | 69–99 | 68–99 | 68–102 | 68–102 | 68–102 | 69–102 | 68–102 | 68–102 |
| **α4** |  | 109–119 | 109–119 | 109–119 | 109–119 | 108–119 | 109–119 | 109–119 | 109–119 | 109–119 |
| **α5** | **TM4 + FHIPEP** | 123–173 | 123–173 | 123–173 | 123–173 | 123–173 | 123–173 | 123–173 | 123–173 | 123–173 |
| **α6** | **TM5 + FHIPEP** | 178–229 | 178–229 | 178–229 | 178–230 | 178–230 | 178–229 | 178–229 | 178–229 | 178–229 |
| **α7** | **TM6** | 235–273 | 235–272 | 235–273 | 235–272 | 235–272 | 235–272 | 235–272 | 235–273 | 235–273 |
| **α8** |  | 277–287 | 277–287 | 277–287 | 277–287 | 277–287 | 277–287 | 277–287 | 277–287 | 277–287 |
| **α9** | **TM7** | 289–304 | 289–304 | 300–304 | 289–304 | 289–304 | 289–304 | 289–304 | 289–304 | 289–304 |
| **α10** | **TM8** | 312–330 | 311–330 | 311–330 | 312–330 | 312–330 | 312–330 | 312–330 | 312–330 | 312–330 |

Extended Data Table 3. CryoEM data collection, processing, and refinement statistics of the wild-type FlhA_C_ ring structure.

| Dataset | FlhA_C_ in C1 symmetry | FlhA_C_ in C9 symmetry | Chemical-crosslinked FlhA_C_ in C1 symmetry | Chemical-crosslinked FlhA_C_ in C9 symmetry |
| --- | --- | --- | --- | --- |
| EMDB  PDB | EMD-80484  25YY | EMD-80485  25YZ | EMD-80486  25ZA | EMD-80487  25ZB |
| **Data collection and processing** | | | | |
| Magnification | 60,000 | 60,000 | 60,000 | 60,000 |
| Voltage (kV) | 300 | 300 | 300 | 300 |
| Electron exposure (e−/Å^2^) | 80 | 80 | 80 | 80 |
| Defocus range (μm) | -1.8 – -2.2 | -1.8 – -2.2 | -1.8 – -2.2 | -1.8 – -2.2 |
| Pixel size (Å) | 0.786 | 0.786 | 0.786 | 0.786 |
| Symmetry imposed | C1 | C9 | C1 | C9 |
| Imported movies (no.) | 17,833 | 17,833 | 11,643 | 11,643 |
| Initial particle images (no.) | 2,256,923 | 2,256,923 | 5,484,473 | 5,484,473 |
| Final particle images (no.) | 292,355 | 274,285 | 447,294 | 447,294 |
| Map resolution (Å) | 3.3 | 3.05 | 2.87 | 2.6 |
| FSC threshold | 0.143 | 0.143 | 0.143 | 0.143 |
| **Refinement** | | | | |
| Initial model used (PDB code) |  | 3A5I |  |  |
| Model resolution (Å) | 3.3/3.3/3.6 | 3.0/3.0/3.2 | 2.8/2.9/3.1 | 2.6/2.6/2.7 |
| FSC threshold | 0/0.143/0.5 | 0/0.143/0.5 | 0/0.143/0.5 | 0/0.143/0.5 |
| Model vs. Data CC (mask) | 0.82 | 0.84 | 0.82 | 0.82 |
| (volume) | 0.82 | 0.83 | 0.82 | 0.82 |
| **Model composition** | | | | |
| Non-hydrogen atoms | 23,895 | 23,895 | 23,895 | 23,895 |
| Protein residues | 3,078 | 3,078 | 3,078 | 3,078 |
| Ligands | 0 | 0 | 0 | 0 |
| **R.m.s. deviations** | | | | |
| Bond length (Å) | 0.003 | 0.003 | 0.003 | 0.003 |
| Bond angles (^ο^) | 0.613 | 0.678 | 0.461 | 0.439 |
| **Validation** | | | | |
| MolProbity score | 1.65 | 1.47 | 1.78 | 1.54 |
| Clash score | 9.24 | 8.73 | 9.56 | 6.43 |
| Rotamer outliers (%) | 0 | 0 | 2.28 | 1.74 |
| **Ramachandran plot** | | | | |
| Favored (%) | 97.12 | 98.24 | 98.30 | 98.24 |
| Allowed (%) | 2.88 | 1.76 | 1.7 | 1.76 |
| Disallowed (%) | 0 | 0 | 0 | 0 |

**Extended Data Table 4. *Salmonella* strains and plasmids used in this study**

| *Salmonella* strain/Plasmid | Relevant characteristics | References |
| --- | --- | --- |
| ***Salmonella*** |  |  |
| SJW1368 | ∆*cheW–flhD* | 1 |
| NH001 | ∆*flhA* | 2 |
| NH003 | ∆*fliH-fliI ∆flhA* | 2 |
| NH004 | ∆*fliH-fliI flhB(P28T)* *∆flhA* | 2 |
| **Plasmids** |  |  |
| pTrc99AFF4 | Modified pTrc expression vector | 3 |
| pTrc99ES | Modified pTrc expression vector | 4 |
| pMM130 | pTrc99AFF4/ FlhA | 5 |
| pMMGN101 | pGEX-6p-1/ GST-FlgN | 6 |
| pMKM130ES | pTrc99ES/ His-FlhA | This study |
| pMKM130-207 | pTrc99AFF4/ FlhA(Δ331–347) | This study |
| pMKM130-207ES | pTrc99ES/ His-FlhA(Δ331–347) | This study |
| pMKM130-212 | pTrc99AFF4/ FlhA(Δ335–347) | This study |

**References**

1. Ohnishi, K., Ohto, Y., Aizawa, S., Macnab, R. M. & Iino, T. FlgD is a scaffolding protein needed for flagellar hook assembly in *Salmonella typhimurium*. *J. Bacteriol.* **176**, 2272–2281 (1994).
2. Hara, N., Namba, K. & Minamino, T. Genetic characterization of conserved charged residues in the bacterial flagellar type III export protein FlhA. *PLOS ONE* **6**, e22417 (2011).
3. Ohnishi, K., Fan, F., Schoenhals, G.J., Kihara, M. & Macnab, R.M. The FliO, FliP, FliQ, and FliR proteins of *Salmonella typhimurium*: putative components for flagellar assembly. *J. Bacteriol.* **179,** 6092–6099 (1997).
4. Kinoshita, M. *et al*. A β-cap on the FliPQR protein-export channel acts as the cap for initial flagellar rod assembly. *Proc. Natl. Acad. Sci. USA* **122**, e2507221122 (2025).
5. Kihara, M., Minamino, T., Yamaguchi, S. & Macnab, R.M. Intergenic suppression between the flagellar MS ring protein FliF of *Salmonella* and FlhA, a membrane component of its export apparatus. *J. Bacteriol.* **183,** 1655–1662 (2001).
6. Minamino, T. *et al*. Interaction of a bacterial flagellar chaperone FlgN with FlhA is required for efficient export of its cognate substrates. *Mol. Microbiol.* **83**, 775–788 (2012).
